# Challenges and advances in measuring phenotypic convergence

**DOI:** 10.1101/2022.10.18.512739

**Authors:** David M. Grossnickle, William H. Brightly, Lucas N. Weaver, Kathryn E. Stanchak, Rachel A. Roston, Spencer K. Pevsner, C. Tristan Stayton, P. David Polly, Chris J. Law

**Affiliations:** Oregon Institute of Technology, Natural Sciences Department; University of Sheffield, School of Biosciences; Kent State University, Department of Earth Sciences; University of Washington, Department of Biology; Seattle Children’s Research Institute; University of Oxford, Department of Earth Sciences; Bucknell University, Department of Biology; Indiana University, Department of Earth and Atmospheric Sciences; University of Texas, Austin, Department of Integrative Biology

## Abstract

Tests of phenotypic convergence can provide evidence of adaptive evolution, and the popularity of such studies has grown in recent years due to the development of novel, quantitative methods for identifying and measuring convergence. These methods include the commonly applied *C*1–*C*4 measures of Stayton (2015), which measure morphological distances between lineages, and Ornstein-Uhlenbeck (OU) evolutionary model-fitting analyses, which test whether lineages convergently evolved toward adaptive peaks. We test the performance of *C*-measures and other convergence measures under various evolutionary scenarios and reveal a critical issue with *C*-measures: they often misidentify divergent lineages as convergent. We address this issue by developing novel convergence measures—*Ct*1–*Ct*4-measures—that measure distances between lineages at specific points in time, minimizing the possibility of misidentifying divergent taxa as convergent. *Ct*-measures are most appropriate when focal lineages are of the same or similar geologic ages (e.g., extant taxa), meaning that the lineages’ evolutionary histories include considerable overlap in time. Beyond *C*-measures, we find that all convergence measures are influenced by the position of focal taxa in phenotypic space, with morphological outliers often statistically more likely to be measured as strongly convergent by chance. Further, we mimic scenarios in which researchers assess convergence using OU models with *a priori* regime assignments (e.g., classifying taxa by ecological traits), and we find that multiple-regime OU models with phenotypically divergent lineages assigned to a shared selective regime often outperform simpler models. This highlights that model support for these multiple-regime OU models should not be assumed to always reflect convergence among focal lineages of a shared regime. Our new *Ct*1–*Ct*4-measures provide researchers with an improved comparative tool, but we emphasize that all available convergence measures are imperfect, and researchers should recognize the limitations of these methods and use multiple lines of evidence when inferring and measuring convergence.

## INTRODUCTION

Phenotypic convergence is commonly associated with adaptive evolution (e.g., Darwin 1859, Losos 2011), but it can also occur stochastically (Stayton 2008) or as a byproduct of shared developmental constraints (Losos 2011, Speed and Arbuckle 2016). Evidence for adaptive convergence is strengthened when the convergent phenotypes are tied to similar ecological or functional roles, and when the magnitude of convergence is greater than expected by chance. Thus, quantitative examinations of phenotypic convergence are important; they assist in 1) identifying convergence and 2) inferring whether evolutionary change is adaptive. Novel methods for identifying and measuring convergence have recently been developed (Mahler et al. 2013, Arbuckle et al. 2014, Ingram and Mahler 2013, Stayton 2015A, Speed and Arbuckle 2017, Castiglione et al. 2019) and are often accompanied by statistical tests for comparing observed convergence to that which is expected by chance. The increased accessibility of quantitative tests for phenotypic convergence has led to a flood of recent studies on that topic (e.g., Friedman et al. 2016, Zelditch et al. 2017, Baliga and Mehta 2018, Da Silva et al. 2018, Arbour and Zanno 2020, Button and Zanno 2020, Grossnickle et al. 2020, Martinez et al. 2020, Serio et al. 2020, Rincon-Sandoval et al. 2020, Spear and Williams 2020, Baumgart et al. 2021, Huie et al. 2021, Rovinsky et al. 2021, Tamagnini et al. 2021, Alfieri et al. 2022, Bennion et al. 2022, Canale et al. 2022, Law 2022).

Phenotypic convergence is often defined as lineages evolving to be more similar to one another than were their ancestors (Losos 2011, Stayton 2015A, Mahler et al. 2017), and we follow this definition here. This definition is pattern-based, and thus is consistent with any number of evolutionary processes or scenarios which might lead to increased phenotypic similarity among evolving lineages, making it an ideal framework for identifying and quantifying convergence (Stayton 2015A, Stayton 2015B). Further, this definition aligns with the underlying operational definitions of convergence used in recent studies that developed convergence measures (Losos 2011, Arbuckle et al. 2014, Stayton 2015A, Mahler et al. 2017, Castiglione et al. 2019). This strict definition of convergence excludes some patterns that some authors consider forms of convergence (e.g., evolution along parallel trajectories). Thus, we reemphasize here that the chosen definition can significantly influence conclusions of convergence studies, including this study. For instance, relying on a relatively broad definition of convergence that permits focal species to have greater morphological variation than their ancestors (e.g., see ‘imperfect convergence’ of Collar et al. 2014) would increase the chance of concluding that focal taxa are convergent. Moreover, finding a pattern of convergence according to our definition can be used to support a hypothesis of lineages evolving in response to the same adaptive peak; ’imperfect convergence’, although possible under such a scenario, will not provide evidence for shared selection in response to the same adaptive peak. For more information on the definition of convergence, see multiple reviews on the topic, including Moore and Willmer (1997), Stayton (2006), Stayton (2008), Stayton (2015A), Stayton (2015B), Pontarotti (2016), Speed and Arbuckle (2017), Mahler et al. (2017), and Bels and Russell (2023).

The *C*1–*C*4 measures (hereafter, ‘*C*-measures’) developed by Stayton (2015A) are an especially popular means of quantifying morphological convergence (e.g., Friedman et al. 2016, Zelditch et al. 2017, Baliga and Mehta 2018, Da Silva et al. 2018, Arbour and Zanno 2020, Button and Zanno 2020, Grossnickle et al. 2020, Martinez et al. 2020, Rincon-Sandoval et al. 2020, Spear and Williams 2020, Baumgart et al. 2021, Huie et al. 2021, Rovinsky et al. 2021, Tamagnini et al. 2021, Alfieri et al. 2022, Bennion et al. 2022, Canale et al. 2022, Law 2022). *C*-measures are calculated using geometric distances in phenotypic space between focal lineages, relying on ancestral reconstructions for morphologies at ancestral nodes. The underlying feature of *C*-measures is the comparison the maximum phenotypic distance between focal lineages at any points in their evolutionary histories (*D*_max_) and the phenotypic distance between phylogenetic tips (*D*tip) (Fig. 1A). *C*1 is the primary *C*-measure and calculated as 1 – (*D*tip/*D*_max_), with the resulting value representing “the proportion of the maximum distance between two lineages that has been ‘closed’ by subsequent evolution” (Stayton 2015A). In our conceptual illustration (Fig. 1A), two focal lineages have convergently evolved such that their tips are 70% closer to each other than their *D*_max_, resulting in a *C*1 score of 0.7.

**Figure 1.**
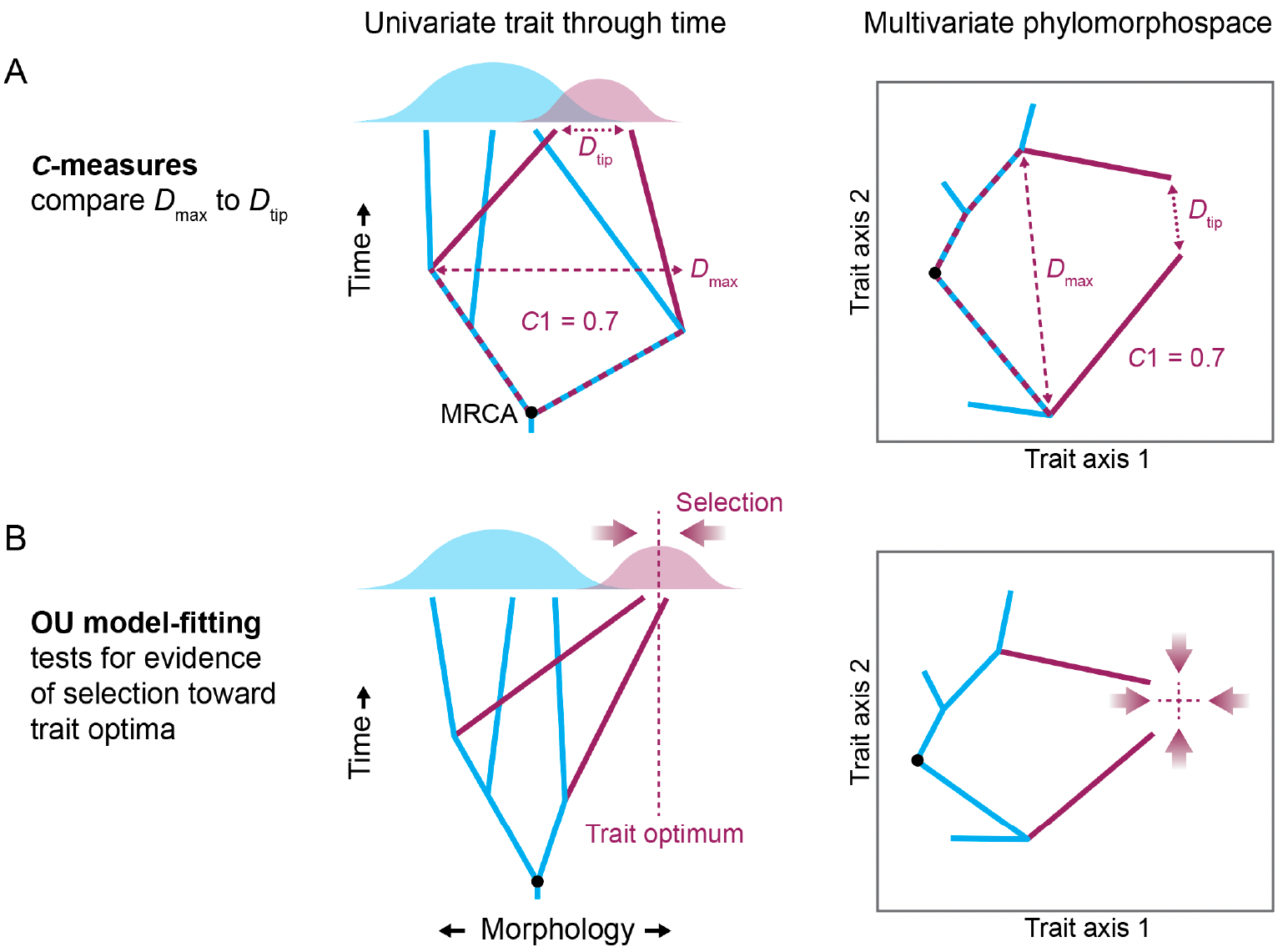
Conceptual illustrations of two methods for assessing phenotypic convergence of focal lineages (maroon): *A*, *C*1 of Stayton (2015A) and, *B*, multiple-regime Ornstein-Uhlenbeck (OU) model-fitting. The *C*1 score of 0.7 indicates that lineages have evolved toward each other to cover 70% of the maximum distance (*D*_max_) between their lineages. *D*_max_ can be measured at any point along the evolutionary histories, starting from the most recent common ancestor (MRCA). *D*tip is the morphological distance between phylogenetic tips. Univariate plots are illustrated as bivariate because there is a time axis, although note that *C*-measure calculations do not incorporate time. *B*, OU models include fitting parameters for trait optima (often interpreted as locations of adaptive peaks) and ‘attraction’ strengths (often interpreted as selection strengths).

One reason for the popularity of *C*-measures is that they can distinguish between convergence (as defined here) and alternative evolutionary patterns that result in distantly related taxa with similar phenotypes, such as conservatism and parallel evolutionary trajectories (referred to as ‘parallelism’ in some contexts). Conservatism is the lack of substantial phenotypic divergence from ancestral morphologies relative to what is expected from various processes (Losos 2008, Moen et al. 2013, McLean et al. 2018); thus, unlike convergence, it does not involve ancestors being less morphologically similar to each other than are their descendants (e.g., see the ‘blue’ lineages in Figure 1B). Similarly, parallelism results in lineages with shared traits, but the lineages’ ancestors never diverged and thus do not reflect convergence as defined here. *C*-measures account for ancestral patterns via the *D*_max_ measurement (Fig. 1A). Alternative distance-based methods for testing for convergence (e.g., Wheatsheaf index, Arbuckle et al. 2014, Arbuckle and Minter 2015; θ, Castiglione et al. 2019) cannot adequately differentiate between convergence and either conservatism or parallelism because phenotypic distances between ancestral morphologies are not considered or, in the case of θ, only partially integrated (Castiglione et al. 2019).

In addition to distance-based measures, evolutionary model-fitting analyses are used to assess convergence, with strong fits of multiple-regime Ornstein-Uhlenbeck (OU) models (Hansen 1997, Butler and King 2004) to morphological data often interpreted as evidence of convergence (e.g., Mahler et al. 2013, Ingram and Mahler 2013, Friedman et al. 2016, Mahler et al. 2017, Baliga and Mehta 2018, Grossnickle et al. 2020, Martinez et al. 2020). An OU process involves ‘attraction’ toward an ‘attractor’ (i.e., trait optimum), which may be due to selective pressures toward adaptive peaks (Fig. 1B). Convergence is sometimes inferred when the best-supported model indicates that two or more focal lineages independently evolved toward the same trait optimum (or adaptive peak). Implementations of these multiple-regime OU models either classify focal taxa into selective regimes *a priori* (e.g., based on a shared ecological trait) or examine whether focal taxa are independently identified (i.e., without *a priori* classifications) as shifting toward a shared adaptive peak. The latter method can be implemented using functions in *R* libraries such as *SURFACE* (Ingram and Mahler 2013; but see Adams and Collyer 2018 for a critique) and *l1ou* (Khabbazian et al. 2016). Other *R* packages can also identify peak shifts (e.g., *Bayou*, Uyeda and Harmon 2014; *phyloEM*, Bastide et al. 2018) but do not include functions specifically designed to identify convergence.

However, an OU process does not require that independent lineages shifting toward a peak (as evidenced by support for OU models) come from more dissimilar ancestors and thus these lineages may not meet the criteria of convergence used in this paper. As such, it may be challenging to use OU model-fitting analyses to differentiate convergence from conservatism; focal taxa may share a given morphology with their common ancestor, but lacking information about ancestral states, it may be difficult to determine whether this is because the taxa retained the ancestral morphology (conservatism) or converged on this ancestral morphology. Nonetheless, a benefit of OU model-fitting analyses is that the magnitude of attraction parameters provides estimates of selective strengths toward adaptive peaks, therefore providing valuable information about underlying evolutionary processes.

We test the performances of *C*-measures and other means of assessing phenotypic convergence using simulated multivariate datasets in which a subset of lineages are modeled as truly convergent or truly divergent. Although our results uncover potential concerns with multiple convergence measures (Wheatsheaf index, θ, and OU model-fitting analyses), the most critical concern is with the *C*-measures: in many circumstances, *C*-measures may lead to misclassification of divergent lineages as convergent. To address this issue, we present an improved method for calculating *C*-measures that minimizes the possibility of erroneously measuring divergent lineages as convergent.

## METHODS

### Evolutionary simulations

We generated simulated trait datasets to ascertain how frequently convergence measures correctly identify *convergent* lineages and misclassify *divergent* lineages as convergent.

Simulated datasets are intended to reflect typical empirical datasets. We simulated traits for 201 species included in a recent phylogenetic treatment of extant mammals, resulting in taxon sampling and phylogenetic structure comparable to recent comparative studies (Grossnickle et al. 2020, Weaver and Grossnickle 2020, Pevsner et al. 2022). The phylogeny is a pruned maximum clade credibility tree produced using 1000 randomly chosen trees from the posterior distribution of Upham et al.’s (2019) ‘completed trees’ analysis. The sample includes 13 gliding-mammal species representing five independent evolutionary origins of gliding behavior. We treated the gliders as the focal lineages (*sensu* Grossnickle et al. 2020); they were the subject of manipulation in our simulations (as such, we refer to those simulated glider data as ’gliders’).

The five ‘glider’ clades are spread across the phylogeny and have varying evolutionary origin ages, making them ideal for representing typical empirical datasets.

Each simulated dataset included six morphological traits, which typically had ranges between 40 and −40 trait units. These are meant to mimic independent linear measurements of an empirical dataset. In the non-glider (i.e., ‘base’) portion of the tree, all six traits were simulated to evolve via Brownian motion (BM). In ‘glider’ clades, between three to six of these traits were simulated to evolve via an OU process (except for drift-based divergence simulations; see descriptions below), with optima varying to produce convergent or divergent trajectories. The BM component of the simulation (‘base simulation’) was performed using the *SimulateContinuousTraitsOnTree* function in the *Phylogenetics for Mathematica* package (Polly 2019). The ancestral value for each trait was arbitrarily set to 0.0 and the step rate (*σ*^2^) was set at 1.0 per million years. In ‘gliders,’ traits evolving via an OU process toward trait optima (see below) were simulated using the *LineageEvolution* function in *Phylogenetics for Mathematica* (ancestral trait values were generated via BM at the base of each clade), with variable optima (θ), a step rate (*σ*^2^) of 0.01, and selection strength (α) of 0.01. See the Supplemental Methods for a description of how these simulations can be replicated using the *OUwie R* library (Beaulieu et al. 2012). Phylogenetic branches of ‘gliders’ were those tipped by one of the 13 ‘glider’ species, plus the subtending branches below clades whose tips were all ‘gliders.’

We generated four types of simulations: 1) convergence of ‘gliders’ via selection (OU-evolved traits), 2) divergence of ‘gliders’ via drift (BM-evolved traits), 3) ‘constrained divergence’ of ‘gliders’ via selection (OU-evolved traits, limited to positive values), and 4) ‘unconstrained divergence’ of focal taxa via selection (OU-evolved traits; no limits on traits). The following text describes these types of simulations, and Figure S1 provides a graphical summary. In total, we generated and analyzed 960 simulated datasets.

*Convergence simulations.* We systematically altered 1) the number of traits (of six total) subjected to convergent selection (three through six) and 2) their optimum values (or ‘targets’) (Fig. S1). Convergent traits were evolved via selection (i.e., OU-evolved) toward the same trait optimum (target). The convergent traits were subjected to directional OU-selection for their full branch lengths, allowing most ‘glider’ tips (besides those with the shortest branches) to arrive at the adaptive peak. Traits (and taxa) not subject to convergent evolution continued to evolve by BM. Note that the OU parameters, apart from the target optimum (i.e., θ), were kept constant (*σ*^2^ = 0.01, α = 0.01) for all simulations involving OU-evolved traits, and variation in the ‘strength’ of convergence was represented by the number of traits subject to convergence rather than the strength of selection per se. Six of six traits being convergent represents extremely strong convergence, whereas three of six traits being convergent represents weaker convergence. Further, we altered the number of convergent traits because empirical datasets often include both convergent and divergent traits (e.g., Grossnickle et al. 2020).

By systematically altering trait optima (‘targets’), we controlled the morphological distance of ‘gliders’ from their ancestral condition, allowing us to test how morphospace position influences results of applied convergence measures (Fig. S1). We used a series of nine trait optima at successively greater distances from the ancestral point in morphospace, starting at zero (convergence toward the ancestral trait values) and successively increasing by 10 trait units to a distance of 80. For example, in a simulation with four traits (of six) being convergent and an optimum of 30, four traits evolved toward an optimum trait value of 30 and the two remaining traits evolved by BM. For simulations using the smallest optima values (zero and 10), the ’gliders’ generally lie within the morphospace occupied by ’non-gliders.’ In contrast, the farthest tested optima (i.e., ∼60–80) result in the ‘gliders’ being very far from ‘non-gliders’ in morphospace. The extreme outliers are unlikely to reflect empirical scenarios but are included to help infer trends. Each simulation with all iterations was repeated with 15 unique ‘base simulations,’ and we report means and standard errors of these 15 replicates.

*Divergence simulation*s. Because divergence may result from both selective and non-selective processes, and because the expected patterns among focal groups varies with these generating processes, we generated two types of divergence simulations. These were drift-based via BM-evolved traits and selection-based via OU-evolved traits. In addition, drift-based simulations involve divergence of lineages that remain relatively closer to the ancestral morphology (i.e., within the morphospace region occupied by non-gliders), which is more difficult to replicate in our OU-evolved divergence simulations (see below). To simulate divergence via drift (i.e., BM-evolved ‘gliders’), we simulated six traits using the *fastBM* function of the *phytools* package (Revell 2012) for *R* (*R* Core Team 2020). For both ‘gliders’ and ‘non-gliders,’ ancestral trait values were set at zero, and, to mimic natural variation, the rate parameter (*σ*^2^) was sampled from a log-normal distribution with log-mean and standard deviation 0 and 0.75, respectively. This was repeated to produce 15 replicate datasets.

For selection-based (OU-evolved) divergence, the five ’glider’ clades each evolved toward a different trait optimum. Between three and six traits were selected toward the clade-specific trait optimum with a series of target distances ranging from 30 trait units from the ancestral morphology (which extends the lineages past the periphery of the base BM tree and thus ensures that the targets are divergent) to 80 units in steps of 10. This means that these divergence simulations are limited to cases in which focal lineages are morphological outliers. A different target was randomly selected for each ’glider’ clade by choosing a random trait value for each of the traits under selection with the condition that their sum of squares equal the squared target distance (i.e., that the target lies at a distance of 30, 40, etc. units from the ancestral trait values). The focal lineages were allowed to fully reach their trait optima.

For our primary selection-based divergence simulations, we ‘constrained’ the OU-evolved ‘glider’ traits to positive values, ensuring that trait optima are divergent yet lie within the same quadrant (or hyper-region) of multidimensional space. This constraint mimics empirical datasets in which lineages exhibit some morphological similarities but are geometrically divergent (e.g., Collar et al. 2014). Nonetheless, we also simulated ‘unconstrained divergence’ (see discussion on subset analyses below) in which the ‘glider’ trait optimum values were not limited to positive values.

Convergence tests applied to simulated data, except some model-fitting analyses, were multivariate, incorporating all six simulated traits. Although our simulated data mimic independent linear measures, all convergence measures tested here could be applied to other data types, including higher-dimensional trait data (e.g., geometric morphometrics data), although long computational times for some analyses might first require data reduction via principal components analyses (PCAs).

### *C*-measures

We applied *C*-measures (Stayton 2015A) to focal lineages (‘gliders’) in the simulated datasets. The primary *C*-measure, *C*1, is calculated as 1 – (*D*tip/*D*_max_), with *D*tip representing the distance between phylogenetic tips of focal taxa and *D*_max_ representing the maximum distance between any tips or ancestral nodes of those lineages (Fig. 1A). The *C*1 value is one for complete convergence and zero for divergence (i.e., *D*_max_ is *D*tip). *C*2 is the difference between *D*_max_ and *D*tip, and it captures the absolute magnitude of convergent change. *C*3 and *C*4 are standardized versions of *C*2, with *C*2 divided by the phenotypic change along branches leading to the focal taxa (*C*3) or the total amount of phenotypic change in the entire clade (*C*4). See Stayton (2015A) for full descriptions of *C*1–*C*4. To calculate *C*-measure scores, we used *R* script functions from Zelditch et al. (2017), which are computationally faster than the functions in v1.0 through v1.3 of the *convevol R* package (Stayton 2015A, Stayton 2018). *C*1–*C*4 scores were calculated for all simulated datasets, but due to computational limits, we only calculated simulation-based *p*-values for a smaller subset of datasets (see the following subsection).

*C*-measures quantify phenotypic convergence between individual tips, not between clades with multiple tips (Stayton 2015A). Thus, we calculated average phenotypes for each focal clade and applied *C*-measures to the five ‘glider’ lineages, each representing an independent evolution of gliding. For example, one glider clade includes six species, so for each simulated trait we calculated mean values for the six species, with the averages used as the representative flying squirrel lineage. These clade averages were only used for *C*-measures.

### Additional measures of convergence

*Subset of simulated datasets*. We applied additional measures of convergence (OU model-fitting, θ, and Wheatsheaf index) to a smaller subset of simulated datasets. This subset only includes datasets in which four of six traits were simulated to converge on a specific trait optimum or diverge toward multiple optima, with the remaining two traits evolved by BM. Four of six traits being convergent results in focal lineages being strongly convergent (e.g., statistically significant distance-based measures of convergence; see Results & Discussion) but not completely convergent. For convergence simulations, we used 15 simulated datasets from sets of simulations where trait optima were set at 0, 20, 40, 60, and 80. These represent simulations in which focal lineages evolve toward the ancestral morphology (optimum = 0), just beyond the morphospace region of BM-evolved lineages (optimum = 20), and far into outlying morphospace (optima = 40, 60, and 80). For selection-based divergence simulations, we used the 15 simulated datasets with optima of 40, 60, and 80. (Using trait optima of 0 or 20 could mistakenly simulate convergence toward ancestral morphologies.) The following methods were only applied to this subset of datasets.

*Evolutionary model-fitting analyses*. We performed evolutionary model-fitting analyses using both multivariate models via functions in the *mvMORPH R* package (Clavel et al. 2015) and univariate models via functions in the *OUwie R* package (Beaulieu et al. 2012). The multivariate models offer the benefit of incorporating all six traits, whereas the univariate models can be set to allow rate (*σ*^2^) and attraction (α) parameters to vary between selective regimes (i.e., ‘gliders’ and ‘non-gliders’) and therefore better represent the underlying simulated data, which include both BM-evolved and OU-evolved lineages.

We fitted four multivariate models to all six simulated traits. The first three models were a single-rate multivariate BM model (mvBM1) that assumes trait variance accumulates constantly and homogeneously on the phylogeny, a single-optimum Ornstein-Uhlenbeck model (mvOU1) that modifies the BM model to constrain each trait to evolve toward a single optimum, and an early burst model (mvEB) that modifies the BM model to decrease evolutionary rate with time. These models do not model ‘gliders’ as evolving to a distinct adaptive peak and thus are not expected to reflect convergence of ‘gliders’ on an adaptive peak (e.g., Fig. 1B). The mvEB model always collapsed to a mvBM model (i.e., the change-in-rate parameter was zero), and thus we did not report mvEB results.

We fitted a two-regime multivariate OU model (mvOU2) that allowed ‘gliders’ and ‘non-gliders’ to exhibit different trait optima (θ). Support for mvOU2 could provide evidence of convergence by indicating that selective forces are driving ‘glider’ lineages to a shared adaptive peak (Fig. 1B). Note that in the simulations ’non-gliders’ evolved via BM (intended to simulate lineages with traits not under strong directional selection or constraint), and thus any support for the mvOU2 model is likely to be driven by the 13 ‘glider’ lineages. Relative support for models was assessed through computation of small-sample corrected Akaike weights (AICcW; Akaike 1974; Hurvich and Tsai 1989). We calculated AICcW for each of the 15 base simulations and report mean values. Prior to fitting the mvOU2 model, we mapped the gliding regime character state onto the ‘glider’ branches of the phylogeny (including the subtending branches below ‘glider’ clades) using the *paintSubTree* function in the *phytools R* package (Revell 2012).

We fitted models to univariate data (PC1 scores) using *OUwie* functions (Beaulieu et al. 2012). Although fitting evolutionary models to PC scores may generate biased results (Uyeda et al. 2015, Adams and Collyer 2018), we fitted these models because, unlike multivariate *mvMORPH* models, they permit evolutionary rates (*σ*^2^) and/or attraction strengths (α) to vary between regimes. We fitted seven univariate models, including single-regime models (BM1 and OU1) and a suite of two-regime OU models (i.e., ‘OUM’ models of Beaulieu et al. 2012). The OU2 model keeps α and σ constant for both ‘glider’ and ‘non-glider’ regimes, OU2A allows α to vary between regimes, OU2V allows *σ*^2^ to vary between regimes, and OU2VA allows both *σ*^2^ and α to vary between regimes (see the Supplemental Methods).

*Additional distance-based convergence measures*. We applied two other convergence measures to the subset of simulated datasets (using all six traits): the Wheatsheaf index, which was implemented via the *R* package *windex* (Arbuckle et al. 2014, Arbuckle and Minter 2015), and θ_real_, which was implemented using the *RRphylo* package (Castiglione et al. 2018, Castiglione et al. 2019). The Wheatsheaf index measures pairwise morphological distances between focal taxa, with distances corrected for the degree of phylogenetic relatedness of lineages. These distances are compared to pairwise distances between other lineages in the sample to determine whether focal lineages are more similar to each other than expected. The θ_real_ measurement (which is different than the θ parameter of OU models) is the angle between the phenotypic vectors of focal lineages (note that these are not vectors of evolutionary change), and it is based upon phylogenetic ridge regression (Castiglione et al. 2018, Castiglione et al. 2019). Smaller angles indicate greater phenotypic similarity, which may reflect convergence of taxa toward a common morphology (Castigione et al. 2019). We report the angle obtained by all pairwise comparisons between focal clades (θ_real_), standardized by the phylogenetic distance separating them (i.e., expected divergence under a BM model).

Significance tests compare standardized θ_real_ values of focal taxa to values computed for randomly selected tip pairs.

## RESULTS & DISCUSSION

### *C*-measure issues

The *C*-measures do not always perform as intended (Stayton 2015A), especially when focal lineages are morphological outliers. This critical problem can manifest in at least two ways (Fig. 2). First, the more outlying the morphologies are in phylomorphospace (and all else being equal, including *D*tip), the greater the *C* scores, indicating stronger convergence. We demonstrate this with a conceptual illustration in Figure 2A. Here, the distances between ancestral nodes and the distances between descendants are unchanged between the scenarios, and thus the *C* scores should be the same for both scenarios. The pattern of greater convergence in outliers is also demonstrated by results of applying *C*-measures to evolutionary simulations (Figs. 3 and S2).

**Figure 2.**
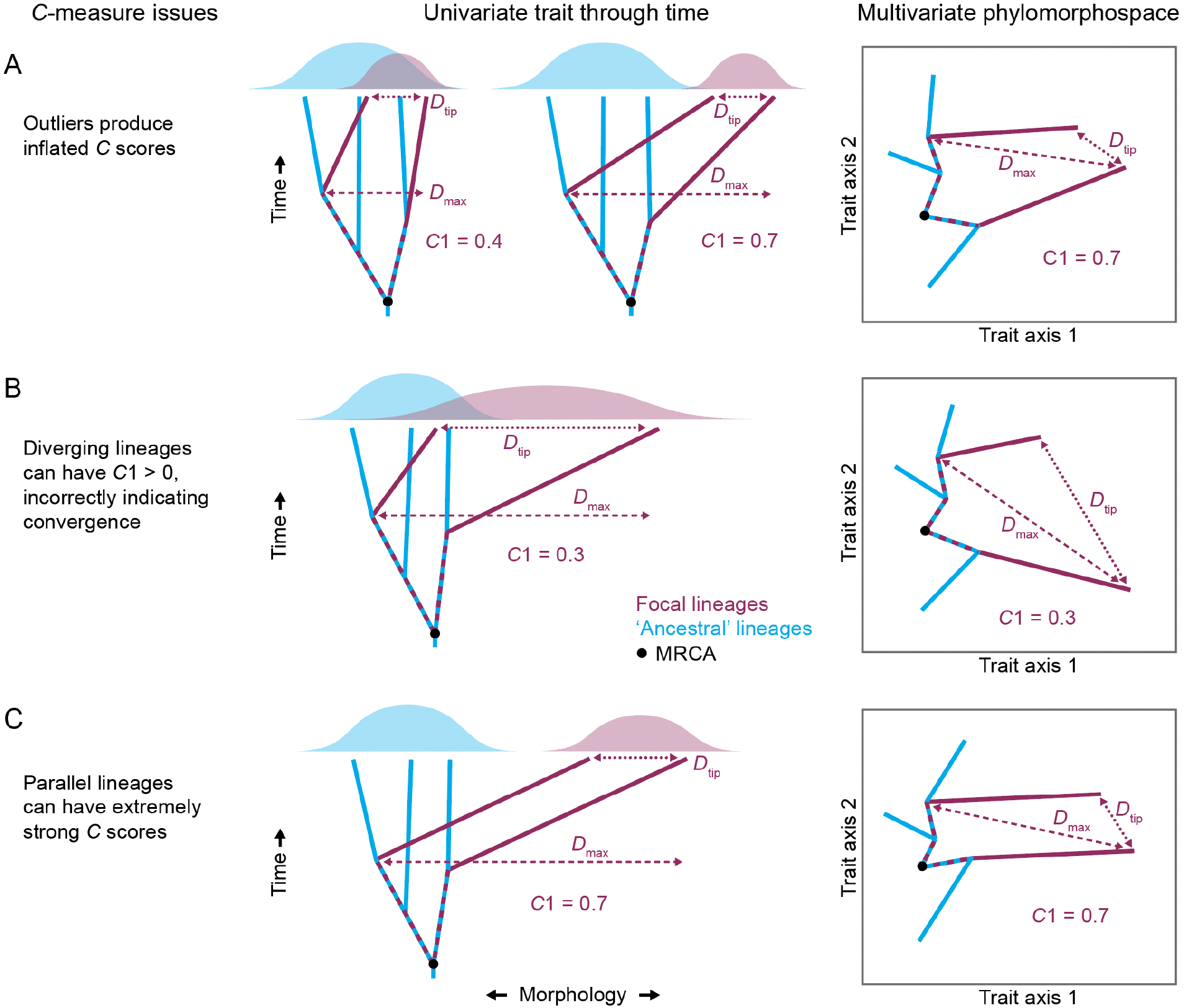
Conceptual illustrations of *C*-measure issues. *C*1 scores are greater than zero for the divergent (*B*) and parallel (*C*) focal lineages (‘maroon’), incorrectly indicating convergence. See the main text and Figure 1 for more information on *C*1, *D*_max_, and *D*tip. The *C*-measure issues may also apply to evolutionary model-fitting analyses because model support for multiple-regime models (often interpreted as evidence for convergence) may be strongest when lineages of different regimes are divergent, such as in *B* (see Results and Discussion).

**Figure 3.**
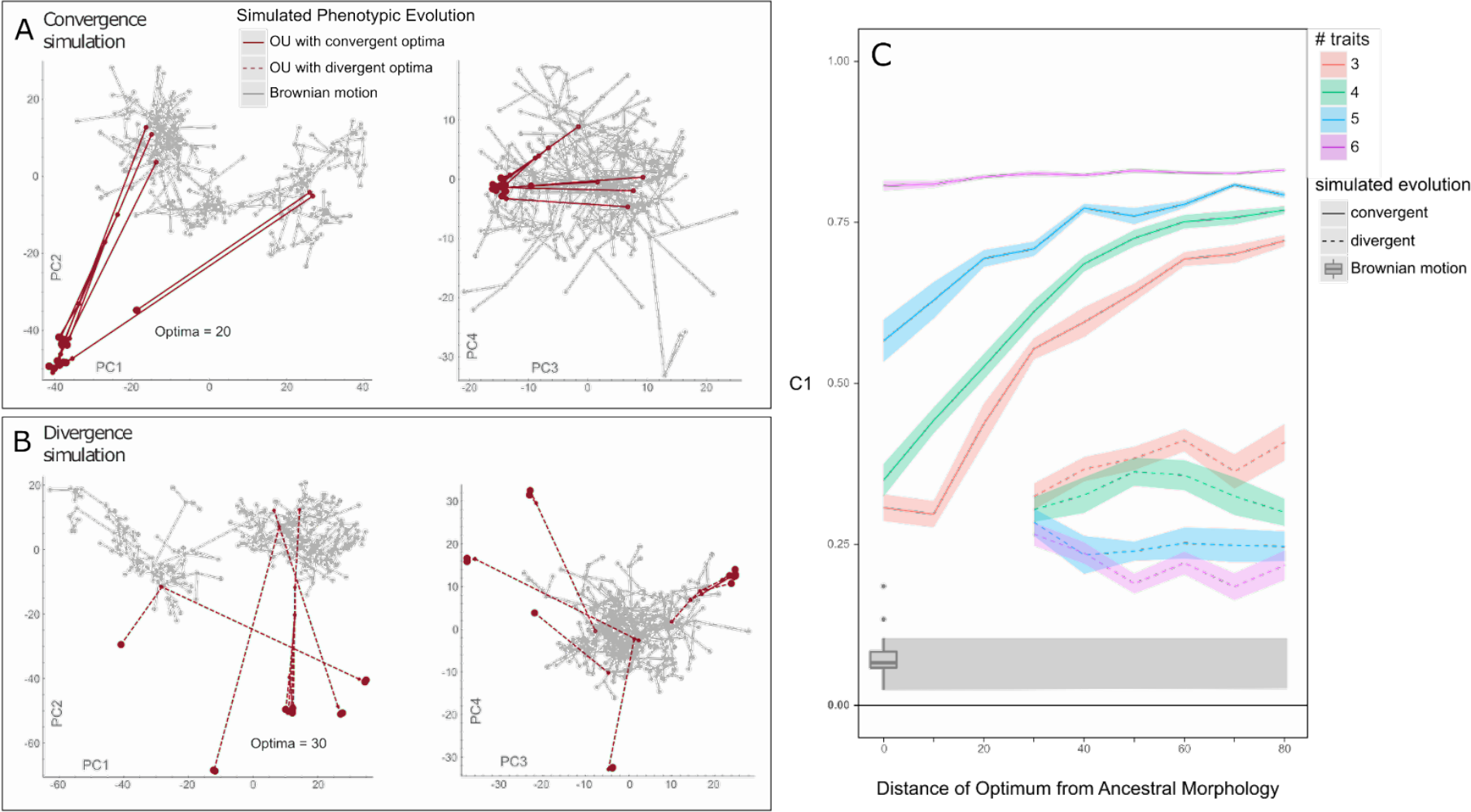
PCA phylomorphospaces for example datasets that simulate convergence (*A*) and divergence (*B*) of all six traits of five focal clades (‘gliders’). After focal clades originated, their traits were selected toward optima via an Ornstein-Uhlenbeck (OU) process, with convergent clades selected toward the same optimum (20 in *A*) and divergent clades selected toward varying optima values (see Methods). (*C*) *C*1 scores (means and standard errors of 15 datasets) for focal lineages, with varying numbers of convergent/divergent traits (of six total) and trait optima positions. Non-convergent/divergent traits and lineages were evolved by Brownian motion (BM). Divergent trait optima are randomized but limited to positive numbers (whereas BM-evolved traits can be negative), resulting in divergent lineages occupying the same region of morphospace (e.g., negative PC2 space in *B*). We did not simulate OU-evolved divergence to trait optima of 0–20 because it often mistakenly generates convergent lineages near the middle of morphospace. But we also simulated divergence via BM evolution, with results as a box-and-whisker plot in *C*.

The second issue with the *C*-measures is more problematic: divergent and parallel lineages can have *C*1 scores greater than zero, incorrectly indicating convergence. In Figure 2B we illustrate lineages that are diverging morphologically but have a *C*1 score (0.3) indicating substantial convergence (closing about 30% of the maximum distance between lineages).

Similarly, lineages evolving along parallel trajectories from a similar ancestral condition (Fig. 2C) have an extremely strong *C*1 score (0.7), which is unexpected because the lineages’ ancestral nodes are the same morphological distance from one another as the distance between tips.

Further, we measured *C*1 in lineages simulated to be divergent (Fig. 3B), and *C*1 values are consistently greater than zero (and statistically significant) for the ‘constrained divergence’ simulations (Fig. 3C; Table 1), incorrectly indicating convergence. *C*1 scores for divergent focal lineages remain around 0.3 at all morphospace positions (Fig. 3B), although the lack of an increase in outliers (like with convergence simulations) may be due in part to methodological choices (see the Supplemental Results and Figure S4).

**Table 1.** Distance-based convergence measures applied to focal lineages (‘gliders’) of simulated convergence and ‘constrained’ divergence datasets (see Methods). Results are means of 15 datasets for each optimum. For θ, we report θ_real_ standardized to phylogenetic distance between clades. Relatively smaller θ_real_ values (i.e., smaller angles between phenotypic vectors) suggest greater convergence, whereas larger Wheatsheaf index, *C*1, and *Ct*1 values indicate greater convergence. Statistical significance (*, *p* ≤ 0.05; **, *p* ≤ 0.01; ***, *p* ≤ 0.001) for *C*1 and *Ct*1 is based on comparisons to 100 BM simulations, for the Wheatsheaf index it is based on bootstrapping with 1000 replicates, and for standardized θ_real_ it is based on bootstrapping with 1000 replicates for each pairwise comparison between the five ‘glider’ clades (except the monospecific clade).

| Simulations | Convergence measure | Trait optimum |  |  |  |  |
| --- | --- | --- | --- | --- | --- | --- |
|  |  | 0 | 20 | 40 | 60 | 80 |
| **Convergence** | $\theta$ (*RRphylo*) | 0.393 | 0.126* | 0.071** | 0.044** | 0.040** |
|  | Wheatsheaf index | 1.976*** | 1.715*** | 1.314 | 0.905 | 0.626 |
| | $C1$ | 0.350* | 0.526*** | 0.685*** | 0.751*** | 0.769*** |
| | $Ct1$ | 0.176* | 0.233** | 0.367*** | 0.497*** | 0.483*** |
| **Divergence ('constrained')** | $\theta$ (*RRphylo*) | – | – | 0.220 | 0.242 | 0.229 |
|  | Wheatsheaf index | – | – | 0.826 | 0.581 | 0.434 |
| | $C1$ | – | – | 0.363** | 0.325** | 0.310** |
| | $Ct1$ | – | – | 0.035 | 0.011 | -0.003 |

The issues with *C*-measures may commonly occur when at least one focal lineage rapidly diverged from other taxa, which can often generate morphological outliers (e.g., see ‘maroon’ lineages in Figure 2B). However, these issues are not limited to outliers and could occur in any region of morphospace. Thus, they have major implications for many empirical studies that rely on *C*-measures (see discussion of examples below); divergent or parallel lineages may often be incorrectly interpreted to be convergent.

The possibility of diverging and parallel lineages having *C*1 scores greater than zero (incorrectly indicating convergence) stems from problematic *D*_max_ measurementss. *D*_max_ can be measured between ancestral nodes (e.g., see Figure 1A), between tips, or between a node and a tip (e.g., see Figure 2). For converging lineages, *D*_max_ is expected to be longer than *D*_tip_ (Stayton 2015A). For diverging lineages, by contrast, *D*_max_ is expected to be the morphological distance between the tips, meaning that *D*_max_ equals *D*_tip_ (and *C*1 = 0). However, this is not always the case; divergent lineages can have a *D*_max_ length that is not between tips, as illustrated in Figure 2B. Thus, divergent lineages can have a *D*_max_ that is greater than *D*_tip_, incorrectly indicating convergence. Although we illustrate this issue using diverging phylogenetic tips (Fig. 2B), the problem could also arise if internal nodes are similarly divergent and outlying in morphospace (and branching lineages from those nodes do not converge on other focal lineages); thus, this issue is not solely due to allowing *D*_max_ to be measured to tips.

### Other measures of convergence

To test the performances of other convergence measures, we applied two ‘distance-based’ metrics (Wheatsheaf index [Arbuckle et al. 2014, Arbuckle and Minter 2015] and θ [Castiglione et al. 2019]) and OU model-fitting analyses to a subset of simulated datasets (Table 1).

*Distance-based convergence measures*. Consistent with the *C*-measures, θ_real_ results (standardized to phylogenetic distance between clades) indicate greater convergence in morphological outliers (Table 1). That is, the angle between phenotypic vectors, θ_real_, decreases when lineages evolve toward optima that are farther from the ancestral morphology. This is unsurprising because all other variables being equal (e.g., phenotypic distance between tips), trait values farther from the origin should result in smaller angles between phenotypic vectors. However, unlike the *C*-measures, θ_real_ does not consistently identify simulated divergent lineages as convergent (Tables 1 and S), although many ‘constrained divergence’ simulations (∼40%) returned significant results. In contrast to *C*-measures and standardized θ_real_, the Wheatsheaf index measures less convergence in outliers relative to non-outliers; values decrease when convergent taxa are farther from the ancestral morphology in morphospace.

*Evolutionary model-fitting analyses*. For datasets with simulated convergent ‘gliders,’ multiple-regime OU models (with a ‘glider’ regime) are the best-fitting models (Table 2), correctly suggesting that ‘gliders’ are converging on a shared adaptive peak (Fig. 1B). However, results are similar when ‘gliders’ are simulated to be divergent – multiple-regime OU models outperform simpler models even though the ‘gliders’ are not evolving toward a shared adaptive peak (Table 2).

**Table 2.**
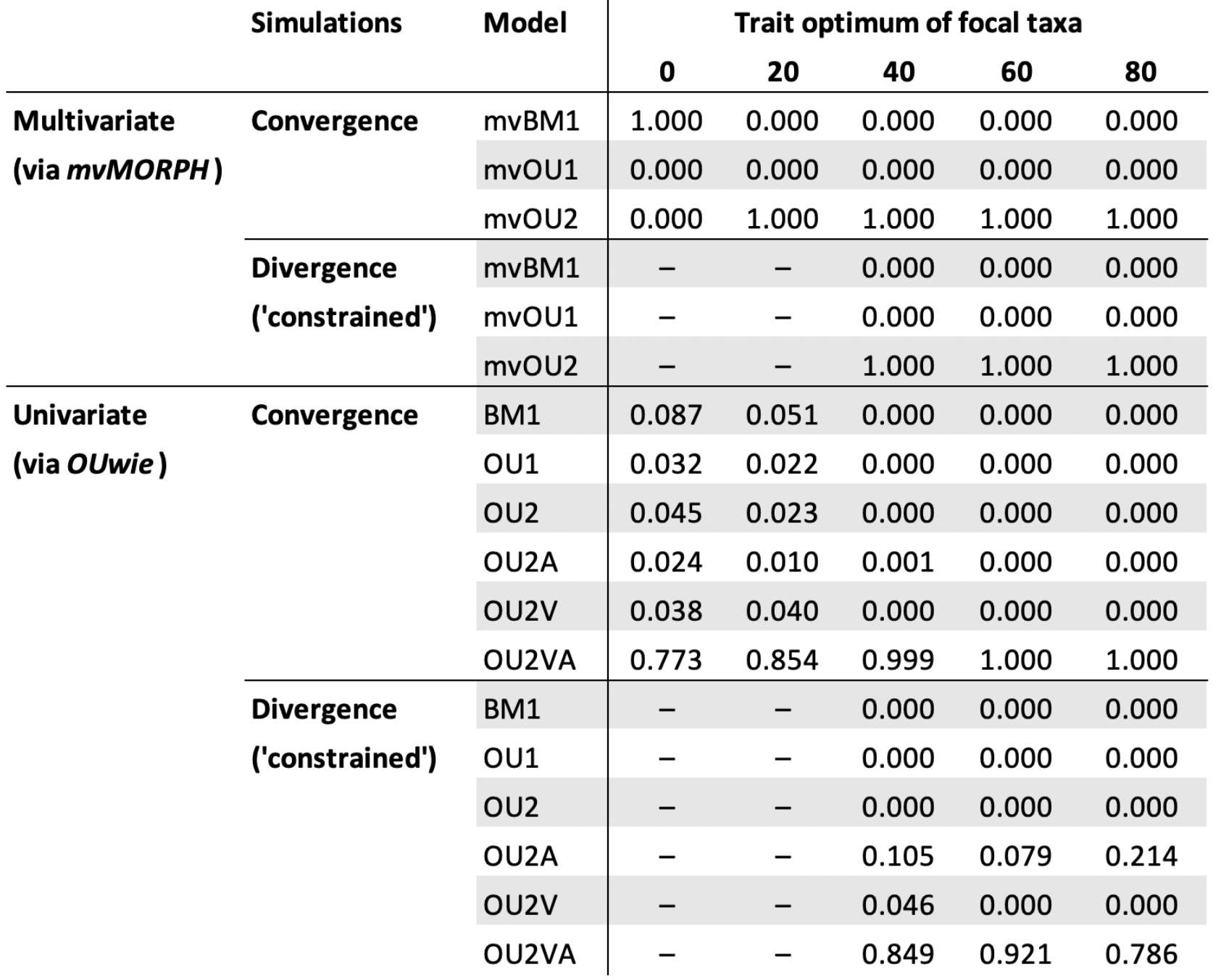
Evolutionary model-fitting analyses in which multivariate models were fitted to all six simulated traits (via *mvMORPH* functions) and univariate models were fitted to PC1 scores (via *OUwie* functions). Results for each trait optimum are mean AICcWs of 15 simulated datasets. Model support for two-regime models (mvOU2 and all OU2 models) could be interpreted as evidence for convergence because these model evolution of focal lineages toward a shared adaptive peak (although see the Results & Discussion for cautionary notes on this assumption). The ‘constrained divergence’ simulations limited the simulated traits of focal taxa (‘gliders’) to be positive values, thus constraining them to one region of morphospace (see Methods). See the Supplemental Results for ‘unconstrained divergence’ results, and see the Methods for descriptions of the models.

Our model-fitting analyses mimic an empirical scenario in which researchers test an *a priori* hypothesis that specific focal taxa (i.e., ‘gliders’; *sensu* Grossnickle et al. 2020) are converging on an adaptive peak. We assigned ‘gliders’ to one selective regime and ‘non-gliders’ to a second regime, and then used model support for multiple-regime OU models as evidence for convergence on adaptive peaks (Fig. 1B). In this type of analysis, support for multiple-regime OU models could suggest that the focal lineages (assigned to a single regime) have convergently shifted toward a shared adaptive peak for that regime. This procedure is prevalent in the literature (e.g., Collar et al. 2014, Friedman et al. 2016, Baliga and Mehta 2018, Grossnickle et al. 2020, Rincon-Sandoval et al. 2020, Law 2022), although other studies assess convergence using model-fitting procedures without *a priori* regime assignments (e.g., Ingram and Mahler 2013, Khabbazian et al. 2016), which are not examined here.

The simulated datasets include lineages evolved by both BM (‘non-gliders’) and a mix of OU and BM processes (‘gliders’; two traits BM-evolved and four traits OU-evolved). Therefore, we do not expect any of the tested multivariate *mvMORPH* models (mvBM1, mvOU1, mvEB, mvOU2) to be good fits because they do not include a mix of BM and OU processes. Although *OUwie* models are expected to provide relatively better fit by allowing a greater number of parameters to vary between regimes, their limitation to only univariate data means they cannot model our full dataset (instead PC1 scores were used).

For convergence simulations, univariate OU model-fitting analyses (i.e., those using *OUwie* models) correctly identify convergence of focal lineages at any optima values (i.e., at any position in morphospace): the OU2VA model is the best-fitting model to PC1 scores and it allows the selection (α) and rate (*σ*^2^) parameters to vary between regimes (Table 2). The results correctly identify different underlying evolutionary processes for the two regimes, with ‘gliders’ showing stronger evidence of convergence via an OU process (i.e., greater α values and smaller *σ*^2^ values than those of non-gliders). This highlights that evaluating parameter values of fitted OU models can assist researchers in evaluating the strength of convergence (and overall fit of the OU model; Boettiger et al. 2012, Ho and Ané 2014, Cooper et al. 2016, Grabowski et al. 2023), with stronger convergence expected to be reflected by relatively larger α values and relatively smaller *σ*^2^ values (e.g., Collar et al. 2014).

However, in multivariate model-fitting analyses (i.e., those using *mvMORPH* models), where rate and attraction parameters cannot vary between regimes, mvBM1 out-performs mvOU2 (suggesting a lack of convergence) when focal lineages converge on a morphology that is similar to the ancestral morphology (i.e., in the middle of morphospace; Table 2). This is expected because there was not a strong shift in morphology from the ancestral morphology, but it highlights that researchers should use caution when applying model-fitting analyses to scenarios in which focal lineages are not morphological outliers. Thus, we recommend including models that allow all parameters to vary among regimes when testing for convergence, because they are more likely to identify convergence when focal lineages converge on ancestral morphologies.

Results for ‘constrained divergence’ simulations highlight a potential pitfall of the assumption that model support for multiple-regime OU models is indicative of convergence (Fig. 1B). Two-regime OU models (both multivariate and univariate) outperformed simpler, single-regime models when fit to data in which lineages of both regimes (‘gliders’ and ‘non-gliders’) are divergent (via ‘constrained divergence’ simulations; Table 2). In these simulations, ’gliders’ do not evolve toward a shared optimum, and are in fact considerably divergent in phylomorphospace (Fig. 3C). Further, ‘non-gliders’ evolved by BM, not an OU process. Thus, taxa of neither selective regime are expected to be well-fit by an OU model, and yet the two-regime OU models are substantially better fits to the data than other models (Table 2). In univariate analyses, the best-fitting model is OU2VA, which allows α and *σ*^2^ to vary between ‘glider’ and ‘non-glider’ regimes – this is the most appropriate model, considering the generating processes of the simulations (i.e., divergent BM and OU processes). However, OU2VA is still a multiple-regime OU model, and its performance could be interpreted as evidence for the presence of two adaptive peaks, one for ‘gliders’ and one for ‘non-gliders,’ even though lineages of each regime were not simulated as evolving toward regime-specific adaptive peaks. This result is consistent with the model-fitting results of Collar et al. (2014) for eel morphology – durophagous eels appear to be divergent in phylomorphospace, but OU models with two regimes (i.e., durophagous and non-durophagous) were the best-fitting models.

That the two-regime OU model is the best fit to divergence data, despite the expectation that none of our fitted models fit the simulated data, may simply reflect that two-regime OU models are the best-fitting of mostly bad-fitting models (with the univariate OUMVA model as a possible exception for some simulated scenarios). Knowing the underlying processes that generated the simulated data, we recognize that the multiple-regime OU models are a poor representation of the data. Nonetheless, we are mimicking specific empirical scenarios in which researchers use *a priori* regime assignments (e.g., based on shared gliding behavior) and do not know the evolutionary processes that generated their trait data. Our results highlight that even if multiple-regime OU models are best-fitting to data, it does not necessarily indicate that the lineages of those regimes experienced convergence (as defined here).

Despite our results, we do not intend to discourage the use of OU model-fitting to test for convergence; they offer a valuable complement or alternative to distance-based metrics. Nonetheless, our results highlight that researchers should be cautious both when choosing models to fit and when interpreting results, especially considering that empirical datasets often include complex evolutionary patterns and small sample sizes, which could influence the fits of OU models (Boettiger et al. 2012, Ho and Ané 2014, Cooper et al. 2016, Grabowski et al. 2023). For robust tests of convergence using an OU-modeling approach, we recommend using models that allow rate (*σ*^2^) and attraction (α) parameters to vary between regimes (e.g., *OUwie* models), with close examination of resulting parameter values. For example, in our univariate model-fitting analyses, although two-regime models had the strongest support in all scenarios, ‘glider’ regimes of divergence simulations have considerably greater *σ*^2^ values compared to those of the ‘glider’ regimes of convergence simulations (Table S2). Thus, in empirical studies, relatively elevated rates be indicative of very weak or no convergence among focal lineages.

We also recommend fitting a broader set of models than those fitted here. For instance, multiple-regime BM models that allow phylogenetic means to vary between regimes, which can be fit using *mvMORPH* functions (Clavel et al. 2015; smean = “FALSE”), may offer more appropriate null models than BM1, and they may complement multiple-regime OU models (e.g., Law et al. 2018, Grossnickle et al. 2020, Martinez et al. 2020, Rincon-Sandoval et al. 2020). Because these models do not model selection toward optima, support over OUM models may suggest limited or no convergence among focal lineages (e.g., Grossnickle et al. 2020). Further, models can be fit that allow for a shift in mode of evolution in focal clades or at a specific point in time (Slater 2013, Clavel et al. 2015), and these may be especially appropriate when different lineages are suspected to have evolved via different evolutionary processes, such as in our simulated datasets, and/or when only one regime is expected to have undergone convergence (reflected by a strong fit of an OU model). If sample size permits, researchers may also consider fitting models that separate focal taxa into multiple regimes. Comparing support for these models, relative to those uniting focal taxa into a shared regime, may aid in identifying false positives like those observed in our simulated divergent datasets.

An additional OU model-fitting approach involves testing for convergent shifts toward shared adaptive peaks without *a priori* regime classifications (e.g., using *SURFACE* functions, Ingram and Mahler 2013; and *l1ou* functions, Khabbazian et al. 2016). We did not apply this method because results are challenging to interpret across many simulated datasets, especially when peak shifts are likely to be identified in BM-evolved taxa (Adams and Collyer 2018), and our approach (using *a priori* regime classifications) may be more appropriate for testing specific hypotheses for focal taxa. Nonetheless, this method provides a valuable research tool, especially for exploratory convergence analyses or as a supplement to other convergence measures (e.g., Rincon-Sandoval et al. 2020).

To mimic empirical samples with focal lineages that share some similarities, our primary divergence simulations (i.e., ‘constrained divergence’) partially constrained ‘gliders’ to one region of morphospace by limiting the randomized optima to be positive values for four divergent traits (the other two traits were BM-evolved and could be negative). This helps to explain the relatively strong fits of the two-regime OU models to the simulated divergence data. Because ‘gliders’ are shifting in similar directions (e.g., all divergent ‘gliders’ are in negative PC2 space in Figure 3B), they may be modeled as evolving toward a broad adaptive peak in one region of morphospace, especially because an OU process does not require that independent lineages shifting toward a peak come from more dissimilar ancestors. Figure 2B provides a conceptual illustration of this scenario; focal lineages (‘maroon’) are diverging but still evolving toward a broad adaptive peak. Further, the eels examined by Collar et al. (2014) may provide an empirical example of this scenario; durophagous eel lineages are morphologically divergent but appear ‘constrained’ to one region of morphospace, and a two-regime OU model is best-fitting to durophagous and non-durophagous eels.

For ‘unconstrained divergence’ simulations, ‘gliders’ diverged in all directions in morphospace, and all convergence measures, including OU model-fitting analyses, correctly identified divergence (Supplemental Results; Table S3). Thus, the central issue for OU model-fitting analyses (i.e., that multiple-regime OU models are best-fitting to divergent lineages) is likely limited to scenarios in which divergent focal taxa share some morphological similarities.

*Summary of issues with convergence measures.* All convergence measures correctly identified convergence among lineages simulated to be convergent (Tables 1 and 2, Figs. 3C and 5A), although results are often influenced by the positions of focal taxa in morphospace. *C*-measures, θ_real_, and OU model-fitting analyses (especially those in which rate and attraction parameters are constant among regimes) show stronger measures of convergence when focal taxa are morphological outliers. In contrast, the Wheatsheaf index shows weaker convergence in outliers (Table 1). For divergence analyses, our most critical finding is that *C*-measures often misidentify divergent lineages as being convergent (Fig. 2B, Fig. 3C, Table 1). In addition, model support for multiple-regime OU models, which is often interpreted as evidence for convergent shifts toward shared adaptive peaks, should be considered with caution because in some scenarios these models may be the best fits to divergent lineages (Table 2).

**Figure 4.**
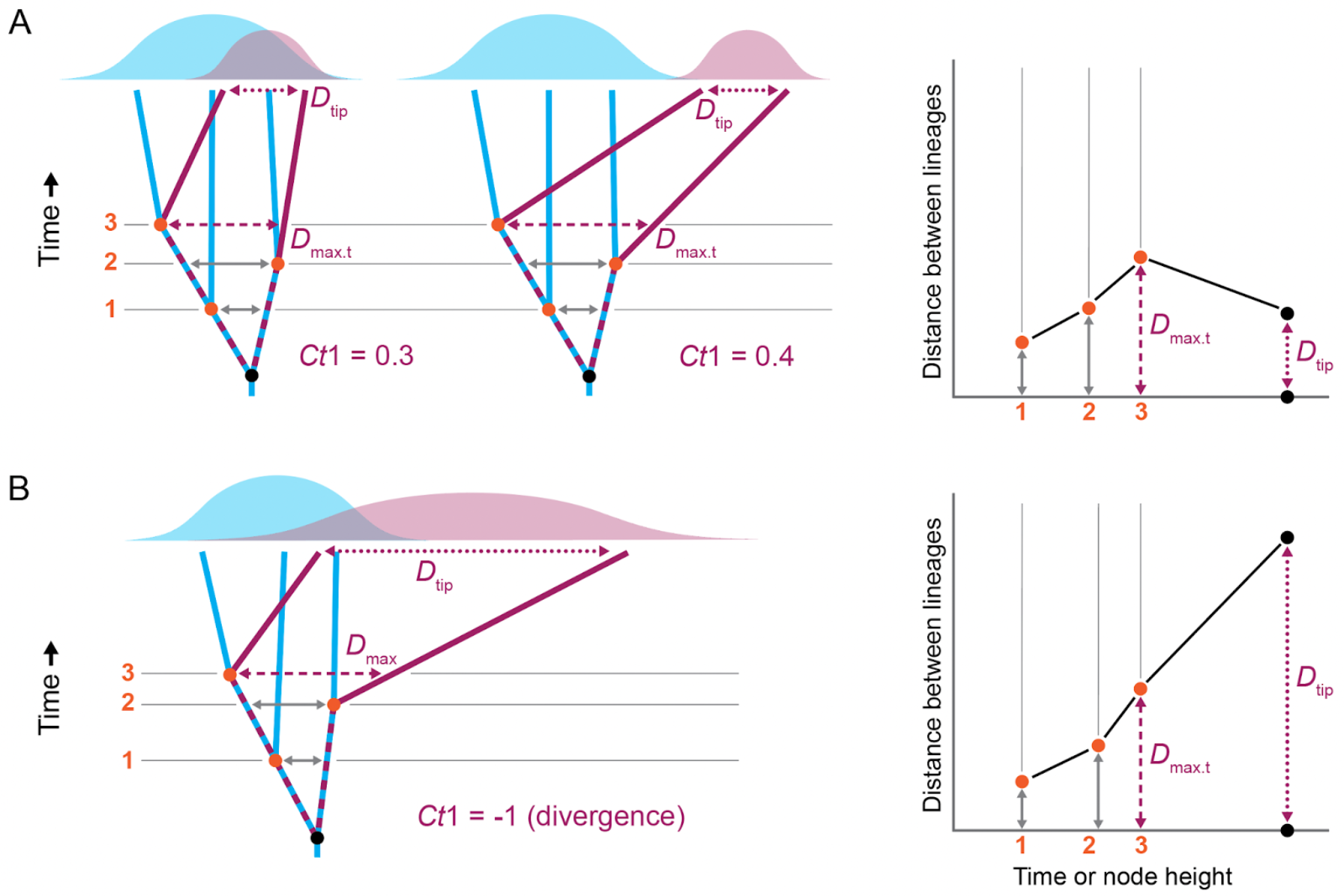
Conceptual illustration of the *Ct*1-measure, which is calculated like *C*1 of Stayton (2015A) but candidate *D*_max.t_ measurements are limited to ‘time slices’ at internal phylogenetic nodes. Plots on the right show the three candidate *D*_max.t_ measurements and the distance between lineages at the tips (*D*_tip_). The scenarios in *A* and *B* are the same as those in Figures 2A and 2B, respectively. In contrast to *C*1, *Ct*1 correctly identifies divergence in *B* (negative score). Although the *Ct*1 score in *A* is greater when lineages are outliers (0.4) versus non-outliers (0.3), the *Ct*1 scores are more similar to each than are the *C*1 scores in the same scenarios (0.7 versus 0.4; Fig. 2A), indicating that *Ct*-measures are less influenced by morphospace positions compared to *C*-measures.

**Figure 5.**
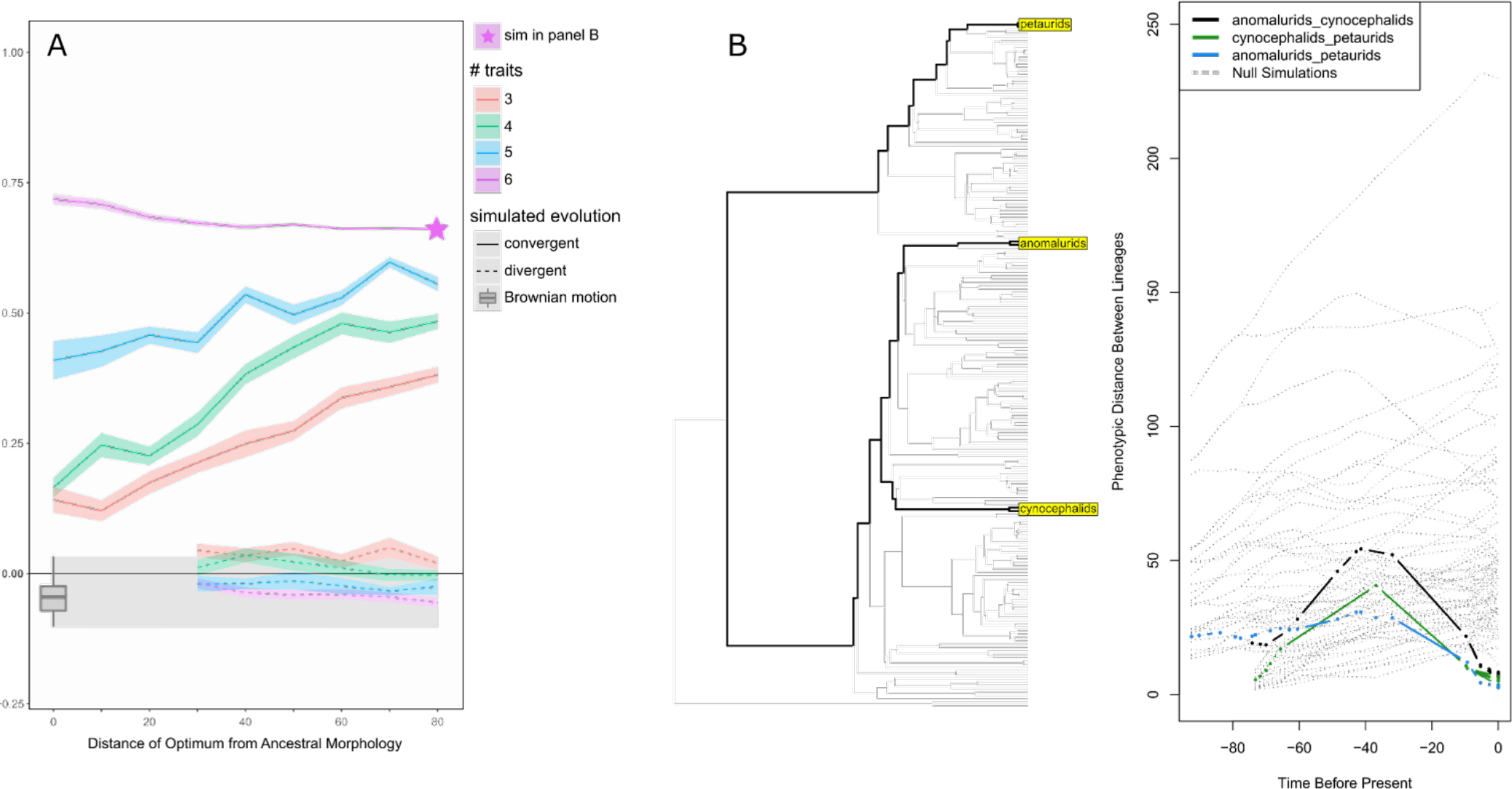
(*A*) *Ct*1 scores for simulated convergent and divergent lineages under varying evolutionary scenarios. See the Methods and Figure 3 caption for more information. Although in some cases the divergence *Ct*1 results are greater than zero (indicating convergence), these results were not statistically significant via simulation-based *p*-values for a subset of datasets (Table 1). *Ct*1 results are means and standard errors of 15 simulated datasets. (*B*) An example output from the *plotCt R* function that shows pairwise distances between lineages with time. Note that although only three ‘glider’ lineages are highlighted in the plot, five lineages were used for *Ct1* measurements.

### Measuring convergence through time via *Ct*-measures

Despite any shortcomings, *C*-measures have benefits over other convergence measures, including being designed to distinguish between convergence and conservatism (Stayton 2015A). Thus, our objective is not to discourage the use of distance-based metrics like *C*-measures but rather to identify issues and encourage the development of improved measures.

We address the *C*-measure issues by presenting novel distance-based convergence measures that are derived from the *C*-measures. The new measures are calculated using the same equations as those for *C*1–*C*4 (except with a change to *C*4; see Supplemental Methods), but we limit candidate *D*_max_ measurements to distances between lineages at synchronous ‘time slices’ coinciding with internal phylogenetic nodes (Fig. 4). We refer to the new measures as *Ct*-measures and *D*_max_ as *D*_max.t_ because time (*t*) is incorporated when measuring morphological distances between lineages. *Ct*1 scores can be interpreted in the same way as *C*1 scores were intended to be interpreted (Stayton 2015A): positive *Ct*1 scores represent a proportion of the maximum morphological distance between lineages that has been closed by convergent evolution, with *Ct*1 = 1 representing complete convergence. Like *C*-measures, statistical significance for *Ct*-measures is based on comparison with expectations for evolution proceeding entirely on a BM model, using simulations to generate the expectations.

By limiting candidate *D*_max.t_ measurements to time slices, *Ct*-measures minimize the possibility of *D*_max.t_ being erroneously inflated by divergent tips. This is conceptually illustrated in Figures 4A and 4B, showing the same scenarios as in Figures 2A and 2B. Whereas the *C*1 score in Figure 2B incorrectly indicates convergence (i.e., *C*1 is greater than zero), the *Ct*1 score in Figure 4B correctly indicates divergence (i.e., the value is negative).

The *D*_tip_ measurement has not been altered from its original implementation in *C*-measures (Stayton 2015A) and is not limited to a synchronous time slice, thus allowing for *D*_tip_ to be measured even when tips vary in geologic age (e.g., an extinct taxon and an extant taxon). However, unlike *C*-measures, the *Ct*-measures do not allow *D*_max.t_ to be measured between tips (i.e., *D*_max.t_ cannot equal *D*_tip_). This results in divergent taxa having negative *Ct* scores (*D*_tip_ is greater than *D*_max.t_), whereas *C*-measures (as initially intended) measure divergent taxa as having scores of zero (*D*_max_ equals *D*_tip_). See the Supplemental Methods for more information on the *Ct*-measures.

The *Ct*-measures include four new features to enhance functionality, which are available via functions (*calcConvCt* and *convSigCt*) in version 2.0.1 of the *convevol R* package (Brightly and Stayton 2023). First, *Ct*-measures can compare clades that contain multiple lineages, whereas the original routines were limited to comparisons of individual lineages (Stayton 2015A). Clade comparisons are enabled by 1) excluding pairwise comparisons between within-clade lineages (e.g., two flying squirrel species) and 2) weighting of *Ct* scores and *p*-values based on the number of pairwise comparisons between focal clades (see Supplemental Methods). Second,

*Ct*-measures can be measured using single traits (the original routines only permitted multivariate analyses, although they were adapted for univariate analyses in some studies; Spear and Williams 2020, Law 2022). Third, users can now specify ancestral states for calculations of *Ct* scores. This is especially useful if a BM model (the default model for inferring ancestral states) is expected to be a poor fit to the data, or if additional constraints should be placed on the traits of interest (e.g., based on fossil evidence). Fourth, we updated the *Ct*4 calculation to better match the original description of that measure. See the Supplemental Methods for additional information on these features.

We used an *R* script written by Jonathan S. Mitchell and published in Zelditch et al. (2017) as a foundation for the updated functions. Run times for the revised *R* functions are approximately ten times faster than the original functions of Stayton (2015A) when using our datasets. We did not revise *C*5, which is a frequency-based convergence measure that tallies the number of times lineages enter a region of morphospace (Stayton 2015A) because it is not influenced by the issues highlighted here.

We have also developed a new *R* function, *plotCt*, that plots pairwise distances between lineages through time. (These distances are also provided as an output of the *calcConvCt* function.) This type of plot is illustrated conceptually (Fig. 4) and in two examples: simulated ‘glider’ lineages (Fig. 5B) and anole ‘twig’ ecomorphotype lineages (Fig. S7; Mahler et al. 2013). These plots allow researchers to visualize when *D*_max.t_ occurred during the evolutionary history of focal lineages and may be useful for applications beyond convergence studies.

We tested the performance of *Ct*-measures by applying them to the simulated datasets using the same methodology as that for *C*-measures. Unlike the *C*-measures (Fig. 3C), the *Ct*-measures do not consistently misidentify divergent lineages as being convergent (Figs. 5A and S3); most of the simulated divergence datasets exhibit *Ct*1 scores that are negative, correctly indicating divergence. Although in some cases the *Ct*1 score is greater than zero when applied to divergence datasets (incorrectly indicating convergence; Fig. 5A), these *Ct*1 scores are not statistically significant, which contrasts with the strongly significant *C*1 scores for divergence simulations (Table 1). Further, the greater-than-zero *Ct*1 results could be due in part to convergence occasionally occurring by chance in our simulated-divergence datasets (e.g., BM-evolved ‘glider’ lineages evolving toward each other).

Although both *C*1 and *Ct*1 have a maximum value of one (complete convergence), the two measures scale differently and thus their values should not be directly compared. For instance, when applied to the ‘constrained’ divergence simulations, *C*1 = ∼0.3 (Fig. 3C) and *Ct*1 = ∼0 (Fig. 5A). Positive *C*1 values reflect both convergence and some degree of divergence (due to the *D*_max_ issue highlighted in this study), and their values are expected to be greater than those of *Ct*1 for the same scenario.

### Relatively greater convergence in morphological outliers

Like *C*-measures and θ_real_, the *Ct*-measure scores are often greater when focal lineages are morphological outliers (Figs. 4A, 5A, and S8). This pattern is more pronounced in *C*-measures compared to *Ct*-measures (Figs. 2A, 3C, and S8), likely due to the *C*-measure issues highlighted in this study (Fig. 2). Further, this trend may occur in *Ct*-measures (despite the *C*-measure-related issues being eliminated) when focal taxa evolve toward outlying regions of phenotypic space, in which case the asynchronous origins of the clades could inflate the *Ct*-measures (see the Supplemental Results and Figure S5 for an explanation of this issue). To help address this issue for *Ct*-measures, we added an optional feature to the *calcConvCt* function that limits candidate *D*_max.t_ measurements to the time prior to the evolution of the focal lineages (e.g., prior to the evolution of the earliest glider clade). We recommend using this option as a supplement to regular *Ct*-measures when focal clades have very different origination ages (see Supplemental Results).

The trend of greater convergence in outliers largely disappears for *C*- and *Ct*-measures when all six simulated traits are OU-evolved to trait optima (i.e., all six traits are convergent) – the *C*1 and *Ct*1 values remain relatively constant at any optimum values (Figs. 3C and 5A).

Conversely, the trend is more distinct when a greater number of traits are BM-evolved. This indicates that the pattern of greater convergence in outliers is influenced by the types of simulated traits (BM-evolved or OU-evolved). We believe that trait type influences this pattern because the relative amount of morphological evolution in OU-evolved traits (vs BM-evolved traits) varies depending on the value of trait optima. For instance, when OU-evolved traits are evolved toward optima near the ancestral morphology (i.e., toward zero), those traits may undergo less morphological change relative to the BM-evolved traits. Thus, there is a relatively greater influence of BM-evolved traits (i.e., divergent traits) on *C*- and *Ct*-measures at smaller optima values, resulting in weaker convergence signals via *C*- and *Ct*-measures. Conversely, OU- evolved traits (i.e., convergent traits) of outliers have undergone relatively greater morphological change, evolving farther from the ancestral morphology, and therefore the outlying OU-evolved traits have a relatively larger influence on *C*- and *Ct*-measures. This pattern likely reflects empirical scenarios because the strength of natural selection is expected to vary among traits (e.g., Grossnickle et al. 2020). Thus, when different traits have undergone varying amounts of adaptive morphological change, researchers should expect to observe relatively stronger evidence of convergence in outliers when using most convergence measures.

### Empirical examples – *C*1 vs *Ct*1

In some cases, erroneous *C*-measure results may have led researchers to infer convergence in lineages that are divergent or infer inflated degrees of convergence. For instance, Grossnickle et al. (2020) tested for convergence among gliding mammal lineages using limb measurements and recovered statistically significant *C*-measure scores, indicated strong convergence. But considering the issues highlighted here, the strong *C*-measure scores in Grossnickle et al. (2020) were likely inflated due to the outlying morphologies of some gliders (e.g., dermopterans) because outliers produce greater *C* scores (Figs. 2A and 3C). This suggests that the gliders are probably less convergent than the authors concluded. We applied *Ct*1 to the data from Grossnickle et al. (2020), and in contrast to strong *C*1 scores, we found that all glider comparisons have negative *Ct*1 scores, indicating divergence instead of convergence. In some instances, *Ct*1 scores are only slightly negative and have significant *p*-values (e.g., *Ct*1 = −0.01 and *p* < 0.01 for the comparison of scaly-tailed squirrels and flying squirrels), which is congruent with other lines of evidence explored in Grossnickle et al. (2020) that suggested glider limbs experienced parallel evolutionary changes rather than convergence.

Huie et al. (2021) and Stayton (2015A; using data from Mahler et al. 2013) independently analyzed *Anolis* lizard morphologies using distinct datasets and found that ecomorphotypes with the greatest *C*1 scores are those in the outermost regions of morphospace (‘crown-giant,’ ‘grass-bush,’ and ‘twig’; see Figure 3 of Huie et al. 2021). The *C*1 values for these ecomorphotypes ranged from 0.31 to 0.43 in these studies, whereas other, non-outlying ecomorphotypes had *C*1 values ranging from 0.09 to 0.25 (Stayton 2015A, Huie et al. 2021). The relatively large *C*1 scores of outlying ecomorphotypes, in addition to the positive *C*1 scores for all pairwise comparisons, may be due in part to the biases in the *C*-measures. We applied *Ct*-measures to one of the outlying ecomorphotypes (’twig’) from the anole dataset of Mahler et al. (2013; ten standardized measurements of body dimensions). We found that, although the overall *Ct*1 score was statistically significant, it was near zero (Table S1), in contrast to the *C*1 score being 0.36 (Stayton 2015A). Interestingly, there was considerable disparity in the pairwise *Ct* results for the five twig lineages, with *Ct*1 scores ranging from 0.346 (*A. paternus* vs. *A. valencienni*) to −0.763 (*A. occultus* vs. *A. paternus*) and six of ten pairwise comparisons not significant. (See Figure S7 and Table S1 for full results and plotted pairwise distances through time.) Thus, these results highlight not only the inflation of *C* scores among outliers, but also the importance of considering individual pairwise comparisons when evaluating convergence among multiple focal lineages.

### *Ct*-measures – recommendations and limitations

In contrast to *C*-measures, the *Ct*-measures are influenced by the timing of evolutionary change because they limit candidate *D*_max.t_ measurements to specific time slices. This feature should be considered when applying *Ct*-measures because it may alter expectations about the degree of measured convergence. For instance, if different focal lineages evolve toward a specific morphology (or adaptive peak) at different points in time, then the *D*_max.t_ measurement may not measure the morphologically farthest distances between the lineages, possibly resulting in lower-than-expected *Ct* scores. Conversely, and as noted above, if focal taxa evolve toward outlying regions of morphospace, then asynchronous origins of clades could inflate *Ct*-measure scores (Supplemental Results; Fig. S5). To help mitigate this issue, we recommend assessing phylomorphospace and distances-between-lineages-through-time plots, and comparing default *Ct* results to those generated when using the alternative option of the *calcConvCt* function that limits candidate *D*_max.t_ measurements to the period in which lineages of interest overlap in time (see Supplemental Methods).

*Ct*-measures can be applied to taxa of varying geologic ages (if there is an internal node for a *D*_max.t_ measurement), but they may perform poorly when tips of focal taxa are very different in geologic age because candidate *D*_max.t_ measurements are restricted to the period of evolutionary overlap among focal lineages. For instance, the evolutionary histories of ichthyosaurs and dolphins overlap from their most recent common ancestor (early amniotes) to the ichthyosaur tips, providing considerable time (more than 200 million years) for candidate *D*_max.t_ measurements. However, there are no candidate *D*_max.t_ measurements between the extinction of ichthyosaurs and the origin of extant dolphins (a period that includes most of placental mammal history). If this is the period in which the ancestral dolphin lineages show the greatest divergence from ichthyosaurs, then *D*_max.t_ will not capture the maximum divergence of the lineages, resulting in smaller-than-expected *Ct* scores. (Note that *D*_tip_ ignores time and would measure the morphological distance between ichthyosaur and dolphin tips.) In these types of cases, alternative methods for measuring convergence (e.g., OU model-fitting analyses) may be more appropriate.

Restricting candidate *D*_max.t_ measurements to coincide with internal nodes exacerbates an issue inherent to many phylogenetic comparative methods: the reliance on inferred ancestral states. *D*_max.t_ is the critical value that enables the *Ct*-measures to diagnose convergence, and it is drawn entirely from ancestral state data, which, by default, are estimated from tip values assuming a BM model of evolution (although users may specify ancestral states obtained by other means). Thus ancestral reconstructions likely reflect average morphologies of the sampled taxa, decreasing the chance of measuring convergence via the *Ct*-measures because *D*_max.t_ estimates may be artificially shorter than the ‘real’ *D*_max.t_ values. We tested this issue by applying *Ct*1 (and *C*1) measures to simulations in which we used the generated (‘true’) ancestral traits, and comparing them to scores obtained using the default BM ancestral state reconstructions and those obtained via alternate methods. *Ct*1 scores are consistently greater when ‘true’ ancestral states are used in calculations (Fig. S8), suggesting the convergence signal of *Ct*-measures might often be diluted due to the assumption of a BM model of evolution. Although different methods for inferring ancestral state values may outperform BM, of those we tested, none consistently reproduced the "true" *Ct*1 scores (Fig. S9). This issue is likely exacerbated when there are relatively few intervening nodes between focal lineages (i.e., there are few candidate *D*_max.t_ measurements), when focal lineages are subtended by long branches (i.e., limiting *D*_max.t_ to deeper nodes), and when using extant-only samples (i.e., there is a lack of fossil data informing ancestral node reconstructions). Therefore, *Ct*-measures may be most appropriate for well-sampled study systems that include a substantial number of internal nodes and relatively few long branches, and researchers should include fossil taxa whenever possible.

The number of phenotypic traits used to assess convergence can influence *Ct* scores. In multivariate datasets, some traits may be convergent and others non-convergent (i.e., divergent, parallel, or conservative). While including a greater number of non-convergent traits in analyses is expected to decrease the overall convergence signal of any convergence measure, it may also exacerbate *Ct*-related issues raised in this section. In general, adding traits increases the measured distances between tips and internal nodes. However, ancestral inference via BM tends to average variation at internal nodes; thus, *D*_tip_ typically increases at a higher rate than *D*_max.t_ for each non-convergent trait that is added to a dataset. This pattern is illustrated in Figure S6, highlighting that increasing the number of simulated BM-evolved traits (expected to be mostly divergent) results in relatively greater increases of *D*_tip_ scores compared to *D*_max.t_ scores. Thus, an increased number of traits in analyses (with all else equal) could result in a relative decrease in *Ct* scores compared to datasets with fewer traits. We recommend choosing traits based on the specific hypothesis being tested (e.g., functional morphological traits that are linked to ecological or behavioral traits of interest) and analyzing individual traits or subsets of traits whenever feasible to tease apart unique patterns among traits.

Given the issues with convergence measures, how should researchers interested in convergence proceed? In general, we believe the newly described *Ct*-measures are the preferred option among distance-based convergence measures, in part because they are largely robust to the biases discussed in this study and can distinguish convergence from conservatism. However, *Ct*-measures may not be ideal in all scenarios, such as when lineages are of very different geologic ages, and alternative methods, including OU model-fitting analyses, may be more appropriate. Further, in cases in which researchers are interested in comparing the degree of similarity among focal taxa, regardless of the mechanism in which lineages evolved similarities (conservatisms, convergence, or parallelism), then alternative geometric methods such as Wheatsheaf index and θ_real_ may be more informative. The OU-approach may be ideal when researchers are interested in the underlying strength of selection that may be driving convergent change, rather than simply measuring convergence. Nonetheless, for OU model-fitting analyses to indicate convergence, the available evidence should go beyond support for a multiple-regime OU model. For instance, researchers should include appropriate null models and examine model parameters for evidence of convergence (e.g., a relatively strong α value is more likely to reflect convergence). Finally, for exploratory analyses of convergence, we recommend researchers implement multiple convergence measures.

### Summary

*C*-measures are a popular means of identifying and quantifying phenotypic convergence, in part because they can differentiate between convergence and conservatism. However, we highlight a critical issue: *C*-measures can misidentify divergent lineages as convergent (Figs. 2 and 3, Table 1). Thus, we developed improved convergence measures (*Ct*-measures) that quantify distances between lineages at time slices at internal phylogenetic nodes, minimizing the possibility of divergent taxa being mistakenly measured as convergent. We also developed new features (available in the updated *convevol R* package; Brightly and Stayton 2023), such as a function to plot distances between lineages through time. Although *Ct*-measures improve on *C*-measures, they have limitations. For instance, *Ct*-measures may be unreliable if convergent evolutionary change is asynchronous between lineages of interest (e.g., fossils of very different geologic ages). More broadly, we find that multiple methods (including *Ct*-measures) are biased by the location of taxa in phenotypic space, with most methods measuring greater convergence in phenotypic outliers. OU model-fitting analyses are a valuable tool for investigating convergence, but we urge caution when applying and interpreting their results. Model support for multiple-regime OU models is not always indicative of convergent shifts toward shared adaptive peaks (Table 2). Because all available convergence measures are imperfect, researchers should consider the benefits and drawbacks of various measures when choosing methods and interpreting results.

## DATA ARCHIVING

Upon publication, all simulated datasets will be archived in Dryad (https://doi.org/XXXXXXX).

## Notes

### Competing Interest Statement

The authors have declared no competing interest.

### Summary of Updates

Edits include shortening of the manuscript, adding clarifications on multiple topics, and adding new supplemental results.

