## Supporting Information for "Challenges and advances in measuring phenotypic convergence"

### SUPPLEMENTAL METHODS

#### Simulations

We produced a graphical summary of our simulations to help clarify the various types and iterations of simulations used in this study (Fig. S1). The four types of simulations include 1) convergence of focal taxa via selection (OU-evolved traits), 2) divergence of focal taxa via drift (BM-evolved traits), 3) 'constrained' divergence of focal taxa via selection (OU-evolved traits, limited to positive values), and 4) 'unconstrained' divergence of focal taxa via selection (OU-evolved traits; no limits on traits). Each set of simulations used six traits per lineage, but for the convergence and 'constrained divergence' simulations the number of traits simulated to be convergent (or divergent) for focal taxa ('gliders') was systematically altered between three and six. Non-convergent traits were evolved via Brownian motion (BM). Varying the number of convergent traits helps to toggle the strength of convergence, with simulations of three convergent traits representing relatively weak convergence. Further, systematically altering the trait optimum values allows us to examine the performance of convergence measures when focal taxa are evolved to different distances from ancestral morphologies (i.e., varying positions in morphospace). See the Methods for more information on each of these types of simulations.

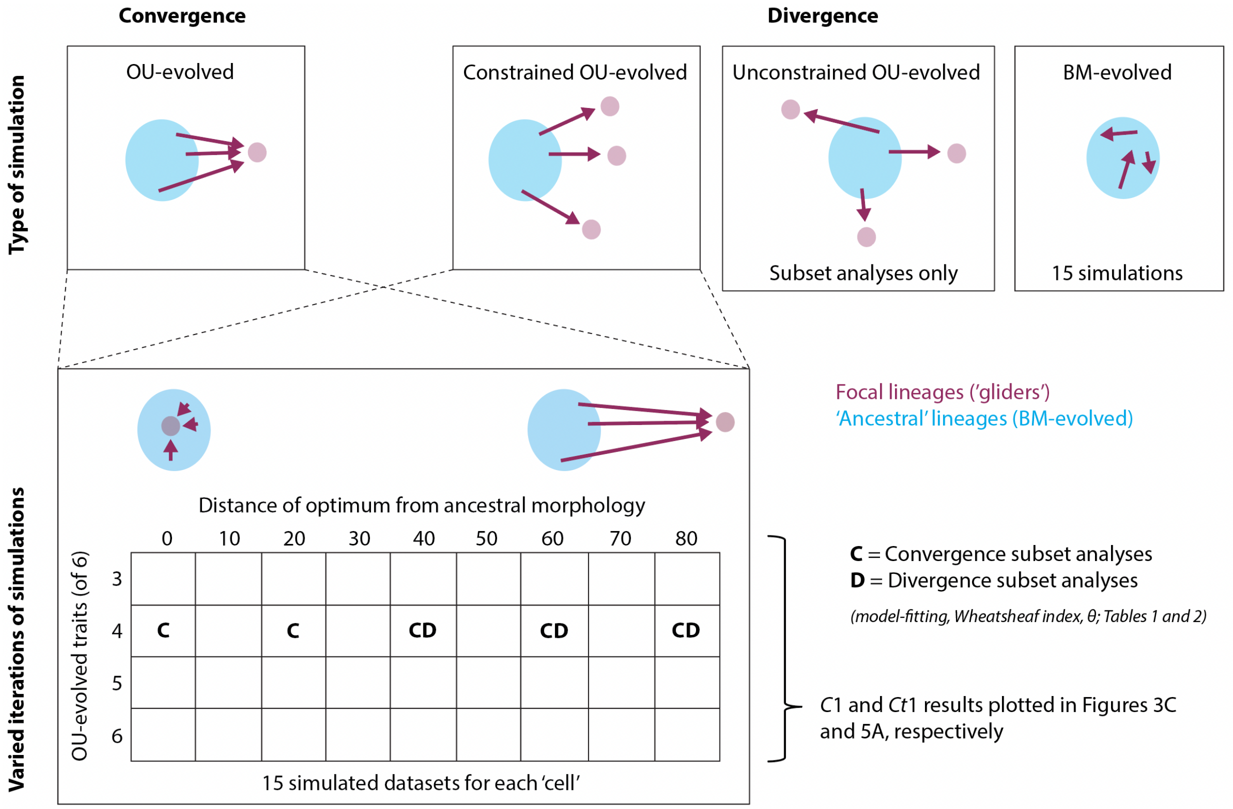


**Figure S1**. A graphical summary of the simulated datasets used in this study. See the Methods for full descriptions. Abbreviations: BM, Brownian motion; OU, Ornstein-Uhlenbeck.

### Univariate model-fitting analyses

A limitation of the *mvMORPH* multivariate models for testing convergence is that they do not permit the evolutionary rate (σ) or strength of attraction to optima (α) to vary between the two selective regimes ('gliders' and 'non-gliders'). This results in poor model performance because the datasets were simulated such that 'gliders' and 'non-gliders' should have very different rates and attraction strengths. For example, the 'non-gliders' are evolved by BM, and thus they are not expected to exhibit attraction to a trait optimum, whereas the convergent 'glider' lineages are expected to exhibit strong attraction due to being simulated by an OU process.

Thus, we also fit seven univariate evolutionary models to the subset of simulated datasets, including several multiple-regime OU models that permit σ and α to vary between regimes. Using functions in the *OUwie R* package (Beaulieu et al. 2012), we fit these models to the first principal component (PC1) scores of a principal components analysis of the six simulated traits. The univariate models include uniform (or single-regime) BM and OU models, as well as a suite of multiple-regime OU models (i.e., 'OUM' models of Beaulieu et al. 2012). The OU2 model keeps α and σ constant for both regimes, the OU2A model allows α (but not σ) to vary between regimes, the OU2V model allows $\sigma^2$ (but not α) to vary between regimes, and the OU2VA model allows both σ and α to vary between regimes.

We recognize that fitting models to PC scores (rather than measurements) can bias results (Uyeda et al. 2015, Adams and Collyer 2018), and thus our univariate results should be considered with caution. However, we feel that using PC1 scores here is justified for two reasons. First, the alternative option is to fit models to each of the six simulated traits individually, but four of the traits are evolved via a strong OU process and two traits are evolved via BM (in our subset of datasets used in model-fitting analyses; see Methods), and thus the model-fitting results are expected to vary considerably between those two types of traits. PC1 provides a single value for which results can be more easily interpreted compared to results for the six traits. Second, the results of the univariate model-fitting analyses are largely congruent with the multivariate results Table 2, with both sets of results showing that two-regime OU models are most commonly the best-fitting models to the convergence and divergence datasets. Thus, the inclusion of the univariate based on PC1 scores does not influence the broad conclusions of this study.

**_Ct_-measures**

We used the *R* script from Zelditch et al. (2017) as a foundation for the updated functions for calculating *Ct*1–*Ct*4 and simulation-based *p*-values because they are computationally faster than the original *R* functions in the *convevol R* package (Stayton 2015, Stayton 2018). The relevant *R* functions are titled *calcConv* (*C* calculations) and *convSig* (significance testing) in the *R* code of Zelditch et al. (2017), *convrat* and *convratsig* in the original *convevol R* package, and *calcConvC* and *convSigCt* for our updated measures.

$D_{max.t}$ *measurement*. The primary change in the *Ct*-measures compared to Stayton's (2015) original *C*-measures is the way in which $D_{\max}$ is defined. *Ct*-measures were designed to ensure $D_{\max}$ (now referred to as $D_{\max.t}$) was obtained from comparisons of synchronous time points along the evolutionary paths leading to the putatively convergent taxa of interest. In this way it prevents the inflation of $D_{max.t}$ that resulted from comparison of asynchronous nodes (e.g., tips and internal nodes) which often occurred when using the original metrics on lineages with outlying morphologies (Figs. 3C and 4). Several modifications to the source *R* code were made to facilitate this change. Candidate $D_{max.t}$ measurements for putatively convergent lineages are now measured at each internal node along the branch paths from the most recent common ancestor (MRCA) of the lineages (e.g., see Figures 4 and 5B). At each of these points we extracted the phenotypic distance between lineages as the euclidean distance between the ancestral reconstruction at the focal node and the coincident reconstruction along the branch path of the other lineage. Where this corresponds to a point along a branch (which is most cases) the ancestral state is estimated using formula [2] from Felsenstein (1985), which allows ancestral states to be interpolated at any point along a given branch from reconstructions at the branch's ancestral and descendant nodes. The code for this was largely repurposed from the *contMap* function of the *phytools R* package (Revell 2012). If no contemporaneous point exists on the opposite path for a given internal node (e.g., when comparing extinct and extant taxa), then a measurement is not taken at that node. All distances measured between paths are stored for each pair of user defined tips. $D_{max.t}$ is the maximum of these distance values, but it is restricted to predate either focal tip (i.e., $D_{max.t}$ cannot equal $D_{tip}$).

Restriction of $D_{max.t}$ to predate the focal tips means the minimum *Ct*1 value is no longer set to zero as in the original *C1*-measure. This allows for some degree of divergence to be captured (i.e., relatively more negative *Ct*1 values may represent greater divergence). However, users are cautioned from using this to test the magnitude of divergence between clades. The *Ct*-measures were not designed to quantify divergence; moreover, in divergent clades $D_{max.t}$ will likely often be the last time point before the oldest focal tip. The method will thus reflect only a small portion of the period when lineages were undergoing divergent evolution. Degree of divergence will then be a function of both phenotypic rates of evolution and of subtending branch length. The latter will in many practical situations be a function of sampling, with long subtending branches due to poor sampling likely to inflate divergence measures substantially since they will provide the best scenario for a large time difference between $D_{max}$ and $D_{tip}$ (and thus capture the greatest proportion of divergent evolution).

The changes to $D_{max}$ were the most consequential of those made to modify the original *C*-measures. However, a number of other new options were also included. These are briefly described below. Full documentation of these options will be available as part of the next update to the *convevol R* package (Stayton, 2018).

*User-defined groups*. The first new option is for users to provide grouping assignments to the tips being tested, thus allowing comparisons of clades with multiple lineages, whereas the original *C*-measures are limited to comparisons of individual lineages. This option removes pairwise comparison between tips within the same group (e.g., two flying squirrels would not be compared if all flying squirrels are defined as one group) and returns results for each unique comparison between groups in addition to overall results. This option is useful if it is hypothesized that two (or more) clades converged, and relieves the user from needing to average tip values of a clade or manually define all of the desired comparisons. When using this option, the overall (for all pairwise comparisons) and comparison-specific *Ct* and *p* values are returned. Overall results are provided as both raw values (means of all pairwise comparisons, excluding within-group comparisons) and weighted values. The latter allows each inter-group comparison to impact the overall average equally, so that larger within group sample sizes don't skew overall results. For instance, if there are three putatively convergent groups (Group A, Group B, and Group C), and Groups A and B both include a single lineage and Group C includes 10 lineages, then there would be 21 total pairwise comparisons among groups (one for A-B, 10 for A-C, and 10 for B-C). Although constituting one third of the unique inter-group comparisons, *Ct* measurements taken from comparison of Groups A and B constitute less than 5% of those used to compute overall (average) *Ct* values. Thus, Groups A and B have a relatively smaller impact than Group C on the overall *Ct* scores and *p*-values. The weighted output scales the *Ct* results (and associated *p*-values) so that each unique inter-group comparison contributes equally to the overall results, whereas the raw overall result simply reports the mean value for all 21 pairwise comparisons. Both weighted and unweighted values are reported in the default output printed by the updated *convSigCt* and *calcConvCt* functions, but we recommend the weighted result be used by default when comparing groups. Nevertheless, the raw result may be preferable in cases in which researchers believe that the more heavily sampled group(s) should have a larger impact on overall results.

Note that it is possible to define groups even when those consist of a single tip. While doing so will not change which pairwise comparisons the model considers, it will provide the user with unique *Ct* scores and *p*-values for each comparison. This can be especially useful when the degree of convergence varies across the lineages of interest (e.g., see the pairwise results for anole species in Figure S7 and Table S1).

*Conservative $D_{max.t}$ option.* When providing user-defined groups, a conservative $D_{max.t}$ option is available that limits candidate $D_{max.t}$ measurements to a time point predating the origination of both focal groups (i.e., the nodes of the MRCAs of each group). This is to prevent $D_{max.t}$ being skewed by an early transition of one lineage toward a shared adaptive optimum that is outlying in morphospace, which can result in inflated *Ct* scores, especially when the origins of the clades are very different in age. This issue is discussed in the Supplemental Results and illustrated in Figure S5. Note that this option is only meaningful when user defined groups are provided. When one of those groups consist of a single lineage the node immediately ancestral to the tip is used. Using this method, long branches can substantially alter inferred $D_{max.t}$ values. We have provided the option to print relevant information about the restrictions put on $D_{max.t}$ when using this method (by setting VERBOSE = TRUE in *calcConvCt*). We strongly suggest that users investigate the impact of using the conservative $D_{max.t}$ option before committing to significance tests.

*Updated Ct4 computation.* In addition to changes to $D_{max.t}$, we also altered the way in which the *C*4-measure is computed. The new version (called *Ct*4) redefines $L_{tot.clade}$, which is the value used to standardize the *C*2 value ($D_{max}$ subtracted by $D_{tip}$) to obtain *C*4. $L_{tot.clade}$ is described by Stayton (2015) as reflecting the total amount of morphological evolution which occurs in the clade originating with the MRCA of two putatively convergent tips. In the original *C*-measures, $L_{tot.clade}$ values were obtained as a sum of the phenotypic distances from all pairwise comparisons between nodes in the clade, but this does not fully account for phylogenetic structure and is heavily influenced by sampling intensity. We have updated this to now be the sum of the phenotypic distances accumulated along each branch in the clade of interest. This change brings *C*4 closer to the original description of the metric.

*Measuring convergence of single traits.* By default, the original *C*-measures do not support investigation of convergence in a single trait (although see Spear and Williams, 2020; Law, 2022). To circumvent this limitation we have added code to the *convrat.t* function which appends an invariant trait (with value zero) to datasets consisting of a single trait. This approach was taken due to ease of integration with existing code, and although crude will provide the same phenotypic distances as would be obtained from the single trait.

*Model output.* Additional changes were made to increase the amount of information returned to the user and facilitate plotting of results. This includes the addition of the novel *plotCt* function, which is described in the 'Measuring convergence through time via *Ct*-measures' section of the main text (with example output in Figure 5B).

**Generating simulated datasets**

To produce simulated trait datasets for this study, we used custom scripts in Wolfram's Mathematica and the *SimulateContinuousTraitsOnTree* function in the *Phylogenetics for Mathematica* package (Polly 2019). However, these simulations can also be generated using other, freely available, programs. For instance, the following *R* code using the *OUwie.sim* function from the *OUwie R* package (Beaulieu et al. 2012) can be used to approximate the simulated datasets used in this study. The values of *nTrt* and *targ*, respectively the number of convergent traits and the value of the target morphology, are the only values that need to be changed to reproduce the full range of simulated datasets (see Figure 5a). The object map is a tree with the glider regimes painted on using the *paintSubTree* function of the *phytools R* package (Revell 2012). This can be reproduced using the tree files published by Upham et al. (2019) – see main text Methods.

```
nTrt <- 4              # number of convergent traits
targ <- 20             # target for convergent traits
alph <- 0.1            # alpha value for convergent traits
sig <- c(1,0.01)       # sigma^2 value for all traits
```

```
OU <- lapply(rep(targ,nTrt), function(x)
OUwie.sim(map,simmap.tree = TRUE, alpha = c(1e-10,alph), sigma.sq = sig, theta0 = 0, theta = c(0,x)))

OU <- do.call(cbind,OU)

BM <- lapply(rep(0,6-nTrt), function(x)
OUwie.sim(map,simmap.tree = TRUE, alpha = c(1e-10,1e-10), sigma.sq = c(sig[1],sig[1]), theta0 = 0,
theta = c(0,x)))

BM <- do.call(cbind,BM)

replica <- merge(OU[,c(1,which(colnames(OU) == "X"))],BM[,c(1,which(colnames(BM) == "X"))], by =
"Genus_species")
rownames(replica) <- replica$Genus_species
replica$Genus_species <- NULL
colnames(replica) <- paste("V",2:(ncol(replica)+1),sep = "")
```

### SUPPLEMENTAL RESULTS

#### *C*1–*C*4 and *Ct*1–*Ct*4 applied to simulated data

In the main text we only present results for *C*1 (Fig. 3C, Table 1) and *Ct*1 (Fig. 5A, Table 1), which were applied to both the simulated convergence datasets and the simulated divergence datasets. However, Stayton (2015) developed four distance-based convergence measures (*C*1–*C*4) and one frequency-based measure (*C*5), with *C*1 being the primary measure, and we altered *C*1-*C*4 to produce the *Ct*1–*Ct*4 measures. Here, we provide full results for *C*1–*C*4 (Fig. S1) and *Ct*1–*Ct*4 (Fig. S2), which are also applied to both the convergence and divergence datasets. See the Methods and Stayton (2015) for descriptions of the four convergence measures, and see the Methods for information on the simulated datasets. Note that the *Ct*4 measure is calculated differently than the *C*4 measure (see Supplemental Methods). For *C*1–*C*4, all results for divergence simulations are greater than zero (Fig. S2), incorrectly indicating convergence, whereas the *Ct*1–*Ct*4 scores for divergence datasets are generally at or below zero (Fig. S3).

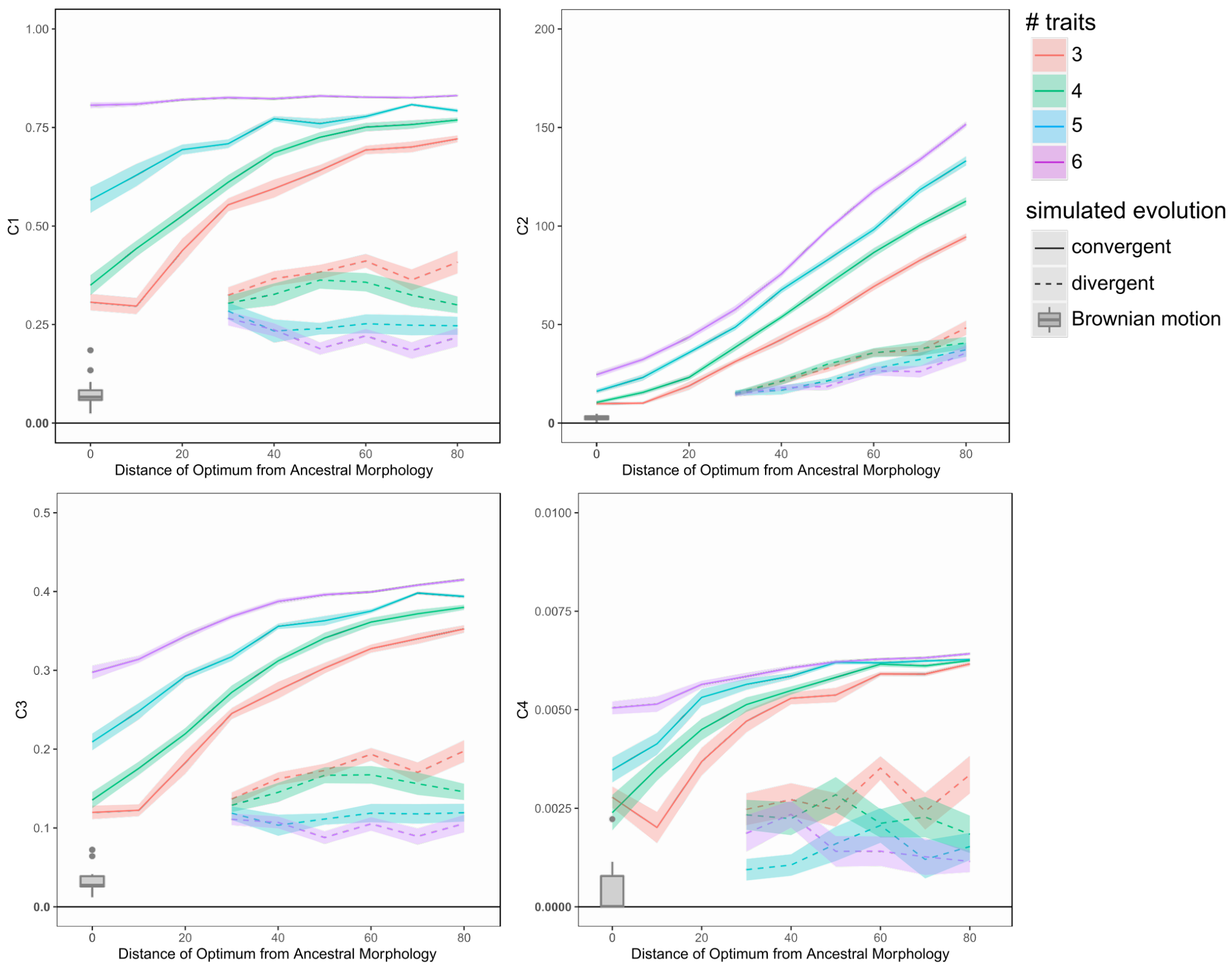


**Figure S2**. Plots of means and standard errors of *C*1–*C*4 scores for simulated convergent lineages (solid lines) and divergent lineages (dashed lines). Datasets varied in the number of convergent/divergent traits (represented by the different colored lines) and in the distance of trait optima from the ancestral morphology (approximated as the center of morphospace). Means and standard errors are computed from 15 simulated datasets. Greater *C*1–*C*4 values indicate greater convergence. We did not simulate divergence for trait optima of 0, 10, and 20 because at these optima our simulation methods may have inadvertently generated convergence patterns (see Methods and Figure 3). As a second means of simulating divergence, we allowed the lineages of interest ('gliders') to evolve via BM. These are provided as box-and-whisker plots, summarizing 15 simulated datasets of six traits (see Methods). Note that the divergence results are all greater than zero, incorrectly indicating convergence.

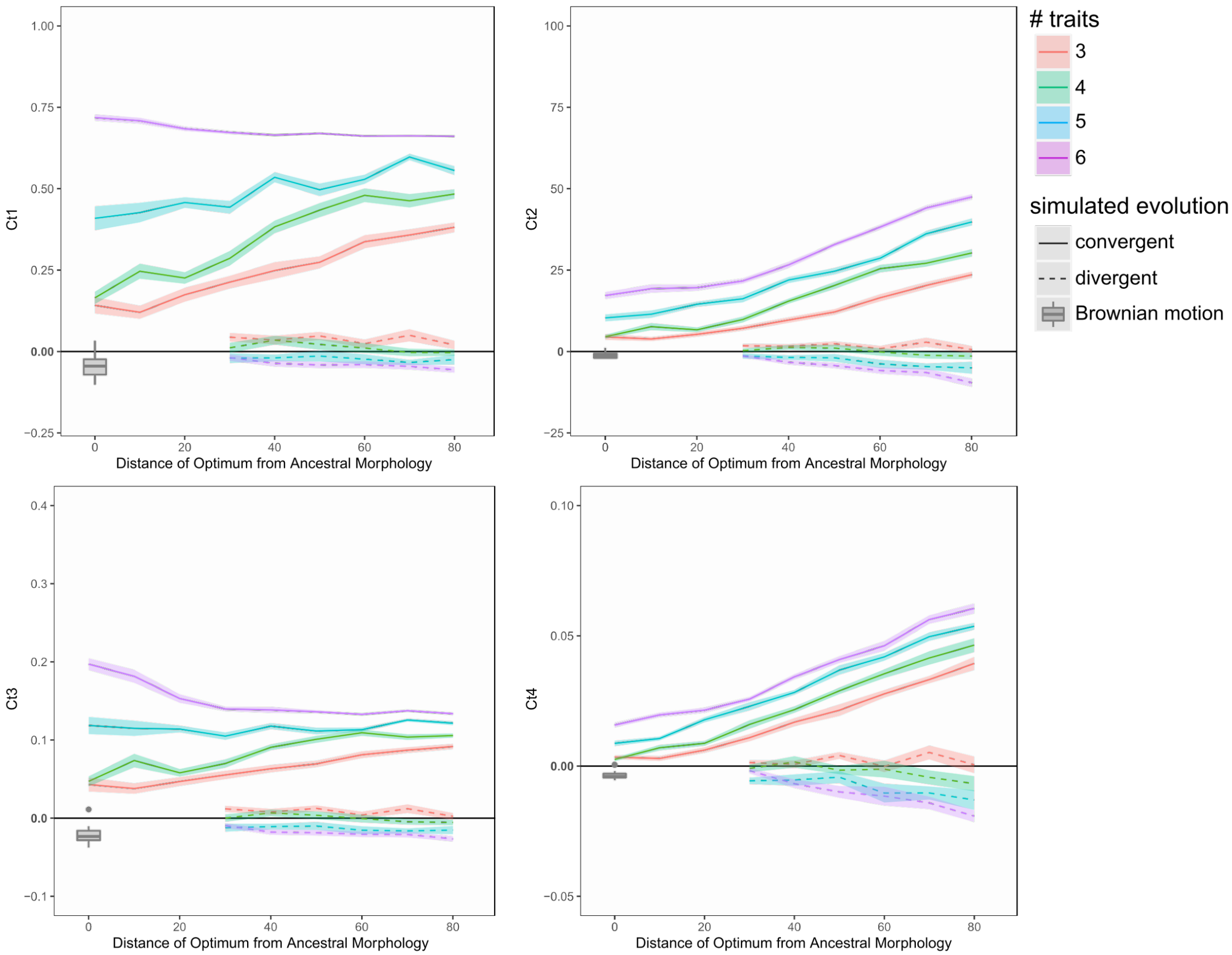


**Figure S3**. Plots of means and standard errors of *Ct*1–*Ct*4 scores for simulated convergent lineages (solid lines) and divergent lineages (dashed lines). Datasets varied in the number of convergent/divergent traits (represented by the different colored lines) and in the distance of trait optima from the ancestral morphology (approximated as the center of morphospace). Means and standard errors are each computed from 15 simulated datasets. Greater *Ct*1–*Ct*4 values indicate greater convergence. We did not simulate divergence for trait optima of 0, 10, and 20 because at these optima our simulation methods may have inadvertently generated convergence patterns (see Methods and Figure 3). As a second means of simulating divergence, we allowed the lineages of interest ('gliders') to evolve via BM. These are provided as box-and-whisker plots, summarizing 15 simulated datasets of six traits (see Methods). Note the differences in the scaling of the vertical axes of the *Ct*2 and *Ct*3 plots relative to the *C*2 and *C*3 plots (Fig. S1), respectively. (The scaling for *C*4 and *Ct*4 is different because these measures are calculated differently.) Also, note the different position of zero relative to results in the *Ct*1–*Ct*4 plots versus the position in *C*1–*C*4 plots (Fig. S1), as well as the overlap in the *Ct*1–*Ct*4 plots of divergence data simulated by both BM and OU processes.

**_C_ and _Ct_ trends for divergence simulations**

For convergence simulations, the *C*- and *Ct*-measures tend to indicate greater convergence involving morphological outliers, at least when some simulated traits are evolved via BM (Figs. 3C and 5A; and see Discussion). In contrast, for divergence simulations the patterns do not show an increase in *C* and *Ct* scores in outliers (Figs. 3C and 5A). This could be interpreted as conflicting with our conclusion that outliers tend to be measured as having greater convergence.

However, the lack of an increase in *C*/*Ct* scores in morphological outliers may be due in large part to a small methodological difference between the convergence simulations and divergence simulations. For convergence simulations, $D_{\text{tip}}$ remains relatively unchanged no matter the position of focal lineages in morphospace. But in contrast, the divergence simulations allow $D_{\text{tip}}$ to increase in lineages that are simulated to evolve toward farther trait optima. This is illustrated in Figure S3, showing that *C*1 remains at ~0.3 when $D_{\text{tip}}$ increases as lineages are greater outliers (compare the left and bottom right illustrations). If we had kept $D_{\text{tip}}$ constant (*sensu* the convergence simulations), then *C*/*Ct* scores would increase in greater outliers (compare the left and top right illustrations in Figure S3). We chose to allow $D_{\text{tip}}$ to vary in divergence simulations because $D_{\text{tip}}$ values may exceed branch length distances between glider species, especially when optima are closer to the ancestral state, and thus it would have been challenging or impossible to keep $D_{\text{tip}}$ constant in all simulations.

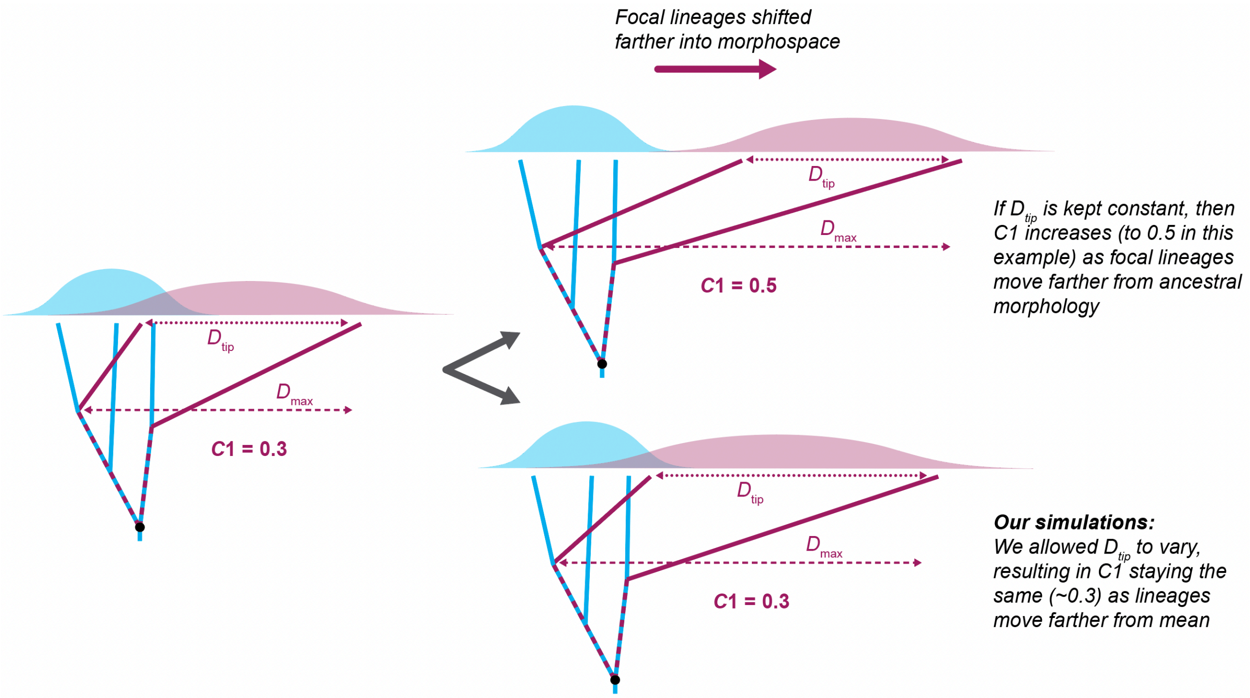


**Figure S4**. Conceptual illustration of how differences in the treatment of simulated divergent lineages can influence the results of *C*- and *Ct*-measures. Unlike the results for convergence simulations, the C1 and Ct1 values do not increase in divergent lineages as they are evolved farther from the ancestral morphology (Figs. 3C and 5A). However, this is likely due in part to $D_{tip}$ increasing in simulations of more outlying divergent lineages (bottom right image).

#### *Ct*-measures – the influence of origination times on results

As discussed in the main text, the *Ct*-measures limit candidate $D_{max.t}$ measurements to specific time slices at internal nodes, and thus the timing of evolutionary change among putatively convergent lineages can influence the results of *Ct*-measures. For instance, if different lineages of interest evolve toward (or away from) a specific morphology at different points in time, then the $D_{max.t}$ measurement may not measure the morphologically farthest distances between the lineages. This issue may be magnified when convergence is expected to be linked to adaptive changes (e.g., adaptations for gliding behavior) that evolved at specific times. For instance, given that colugos (i.e., Dermoptera or ‘flying lemurs’) evolved traits associated with gliding behavior approximately 60 Ma, and flying squirrels (Pteromyini) evolved traits associated with gliding approximately 25 Ma (e.g., Grossnickle et al. 2020), then most of the candidate $D_{max.t}$

measurements will be comparisons of dermopterans with gliding traits to stem flying squirrels without gliding traits (from 60 to 25 Ma). If the older lineage (colugos) has already undergone considerable evolutionary change by the time that the younger lineage (flying squirrels) originated, then much of the convergent evolutionary change of the older lineage is not captured by the morphological distances measured at 'time slices,' which are limited to the time period in which the lineages overlap. Ideally, most candidate $D_{max.t}$ measurements would be comparisons of non-gliding stem colugos and non-gliding stem flying squirrels that lack the adaptive traits associated with gliding. This issue might lead to candidate $D_{max.t}$ measurements being smaller than expected, or at least smaller than those calculated by measures that ignore time (e.g., *C*-measures).

Conversely, if the putatively convergent taxa evolve toward outlying regions of morphospace, then the asynchronous origins of the clades could inflate the *Ct*-measures. We illustrate this in Figure S5. Note that the orange arrows, which represent the origins of the focal lineages (and positions of candidate $D_{max.t}$ measurements), are the same morphological (x-axis) distance apart in each panel if time (the y-axis) is ignored. In the conceptual illustrations, the *Ct*1 score is consistently 0.3 when convergent lineages originate at the same time and/or when lineages evolve toward the ancestral morphology. However, when lineages originate at different times and evolve toward an outlying region of morphospace, then the *Ct*1 score is 0.7 due to an inflated $D_{max.t}$ value (bottom right panel). Thus, researchers should be cautious when applying *Ct*-measures to datasets with outlying taxa of various origination ages, and we offer some suggestions in the main text for mitigating this issue. It is also worth noting that this latter scenario assumes that the convergent lineages can reach adaptive zones; if the later-evolving convergent lineage is still evolving toward outlying morphospace (i.e., it has yet to reach an adaptive peak or zone) then the aforementioned issue may have less of an influence on *Ct* results.

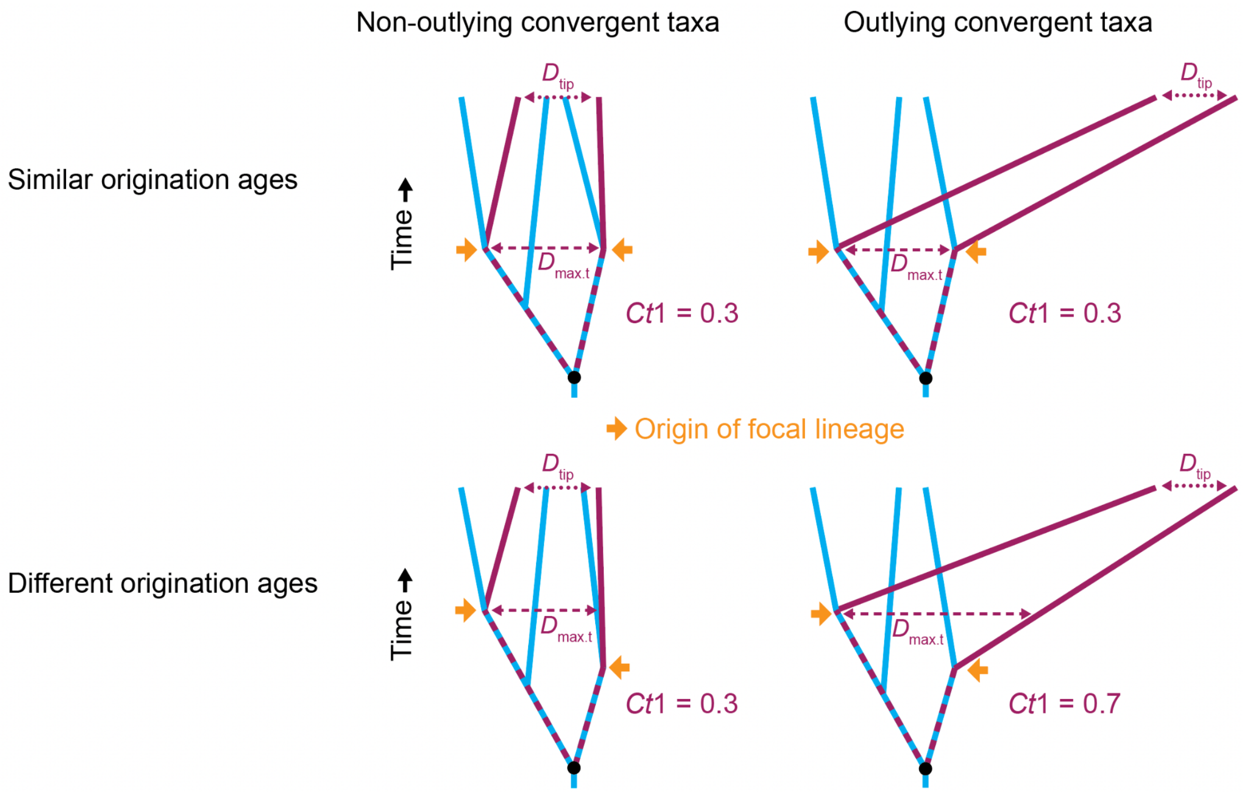


**Figure S5**. Conceptual illustrations demonstrating how *Ct*1 scores can be influenced by a combination of outlying morphologies and varying origination times among convergent focal lineages (maroon). The *Ct*1 score is 0.3 in three of the scenarios but inflates to 0.7 when focal lineages both originate at different times and are outliers in morphospace (bottom right). To help mitigate this issue, we included an option as part of the *convrat.t* function that allows users to limit candidate $D_{max.t}$ measurement to the time period prior to the origination of the focal lineages (Supplemental Methods). See the main text for descriptions of *Ct*1, $D_{max.t}$, and $D_{tip}$.

**Influence of the number of traits on *Ct* results**

As discussed in the main text (see Results & Discussion), the number of traits used in analyses (with all else equal) can bias the *Ct* scores. Inference of ancestral states via BM tends to average variation at internal nodes; thus, $D_{tip}$ typically increases at a higher rate than $D_{max.t}$ for each non-convergent trait that is added to a dataset. (Here, we use "non-convergent traits" to refer to BM-evolved traits that are not selected to evolve toward a trait optimum via an OU process. These are often divergent, although it should be noted that BM-evolved traits could still be convergent by chance.) This is illustrated in Figure S6. The effect of this pattern is that an increased number of traits in analyses (with all else equal) could result in a relative decrease in *Ct* scores, unless those added traits are strongly convergent.

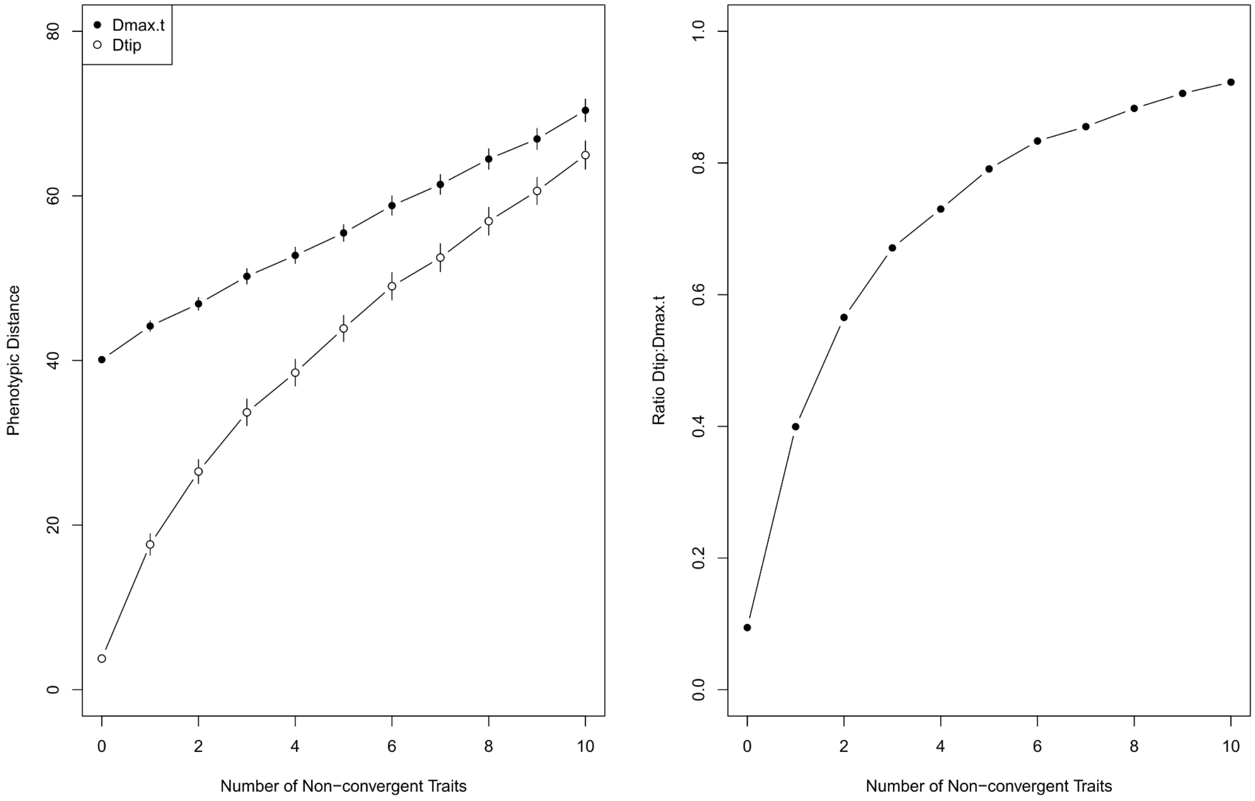


**Figure S6**. Illustration of how the number of traits used in analyses can influence *Ct*-measures, demonstrating the increased rate at which $D_{tip}$ values increase relative to $D_{max.t}$ as additional non-convergent traits are included in analyses. (Here, 'non-convergent traits' refers to BM-evolved traits, which are expected to be divergent in most cases.) The left panel shows $D_{tip}$ and $D_{max.t}$ measured between two 'glider' lineages with two simulated convergent traits (optimum = 80) and varying number of additional traits simulated via BM. Values are averages from 100 simulated datasets with standard error. The right panel shows the ratio between the mean $D_{tip}$ and $D_{max.t}$ values in the left panel.

**Empirical example - *Anolis* 'twig' ecomorphotype**

To test the novel *Ct*-measures and compare *Ct* results to those of *C*-measures (see the *Empirical examples* subsection of the Results & Discussion), we re-analyzed a classic example of convergence among *Anolis* lizards (Mahler et al. 2013), focusing specifically on five 'twig' ecomorphotype lineages. We chose this ecomorphotype because the taxa are morphological outliers that occupy a unique region of *Anolis* morphospace (Huie et al. 2021), and they have especially strong *C*-measure scores (Stayton 2015, Huie et al. 2021), although we believe that this is due in part to the lineages being morphological outliers (see Results & Discussion). Following the methods of Mahler et al. (2013), we size-corrected the traits via PGLS regression of each trait against the snout-to-vent length via PGLS. The *Ct*-measure results for this analysis are provided in Figure S7 and Table S1. Whereas the *C*1 score is 0.36 (Stayton 2015), we find the overall *Ct*1 score to be near zero for both the raw and weighted results (Table S1). This helps to highlight the inflated *C*-measure results due to the issues highlighted in the Results & Discussion. However, note that there is considerable diversity in the results among the ten pairwise comparisons; four are strongly statistically significant, whereas some (e.g., *Anolis occultus* and the *A. paternus* clade) show considerable divergence (*Ct*1 = -0.763; Table S1). To highlight the differences between convergent and non-convergent (or not significant convergence) pairwise comparisons, we separate those comparisons in Figure S7. Thus, we recommend that researchers examine and report results for pairwise comparisons whenever examining more than two putatively convergent lineages.

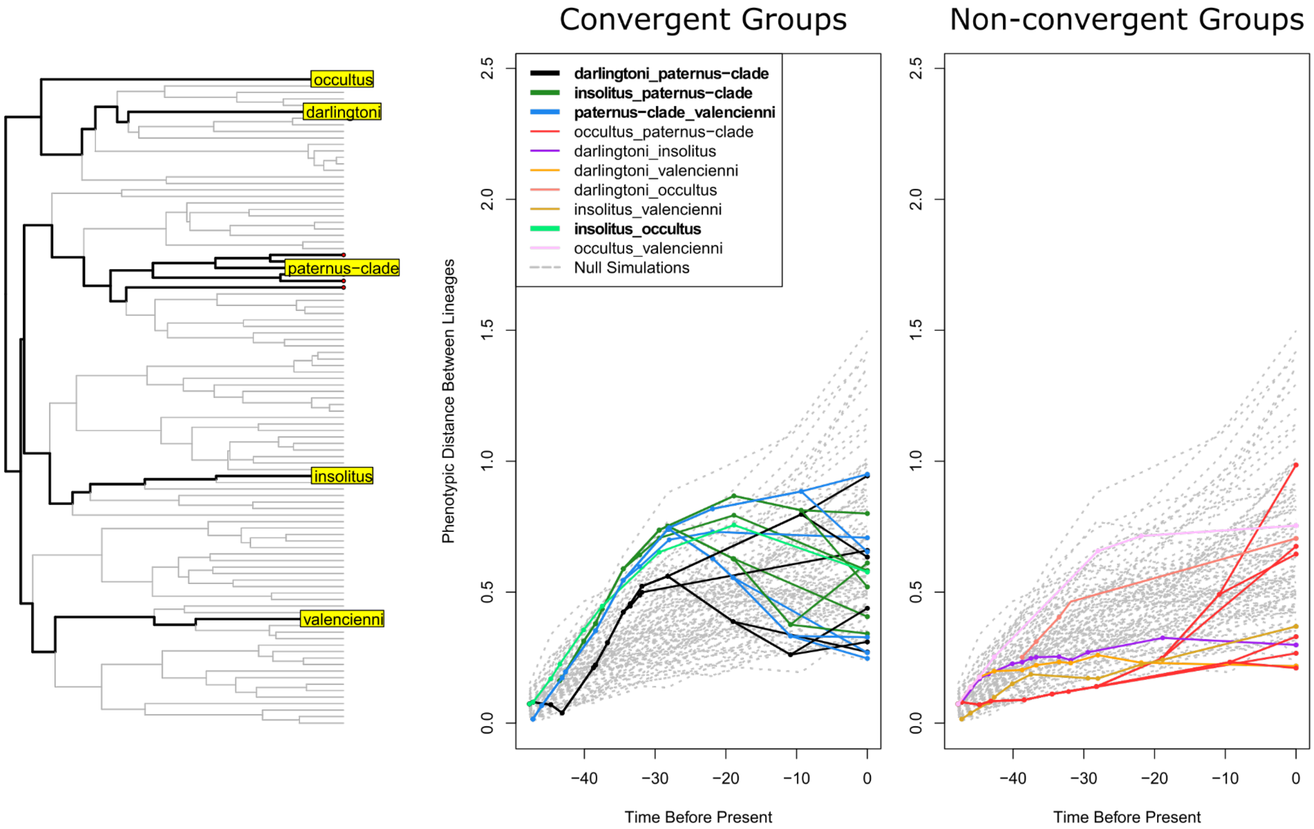


**Figure S7**. Summary of empirical tests of convergence in *Anolis* species belonging to the 'twig ecomorph' (Mahler et al. 2013). We size-corrected (via PGLS regression) and then analyzed the ten skeletal traits of the dataset of Mahler et al. (2013), with taxa assigned to groups based upon unique origins of the 'twig' ecomorphotype (see the *Ct-measures* section of the Supplemental Methods). The plots are the output of the *plotCt* function of the convevol R package, although the distance-through-time plot has been split to show statistically significant (left) and not significant (right) pairwise comparisons separately (see also Table S1). Significant pairwise comparisons are also indicated in bold in the key. Note that two of the 'non-convergent' comparisons in the right panel do have a positive *Ct*1 value, but they are statistically not significant (Table S1). There are 50 null simulations (light gray lines).

**Table S1**. *Ct*-measure values obtained for analyses run using the anole dataset of Mahler et al. (2013; ten standardized skeletal traits). Values are reported for overall comparison of ten 'twig ecomorph' species in five groups (corresponding to each independent origin of the ecomorph; Fig. S7). Pairwise comparisons of groups are also illustrated in (Fig. S7). See the Supplemental Methods for an explanation of the difference between 'overall raw' and 'overall weighted' results. Note that 'pat' refers to a five-species clade that includes *Anolis paternus* and four closely related species, whereas all other 'twig' taxa include a single lineage (Fig. S7); see the Methods for updates to the *convevol R* package that allow for comparisons among taxa with more than one lineage. Asterisks denote values returned as significantly different from null simulations (. - p < 0.1, * - p < 0.05, ** - p < 0.01). Abbreviations: *dar, Anolis darlingtoni*; *ins, Anolis insolitus*; *occ, Anolis occultus*; *pat, Anolis paternus*; *val*, *Anolis valencienni*.

| | Overall | | Pairwise comparisons | | | | | | | | | |
|---|---|---|---|---|---|---|---|---|---|---|---|---|
| | Raw | Weighted | *dar - pat* | *ins - pat* | *pat - val* | *occ - pat* | *dar - ins* | *dar - val* | *dar - occ* | *ins - val* | *ins - occ* | *occ - val* |
| ***Ct* 1** | -0.01** | -0.057** | 0.147** | 0.323** | 0.346** | -0.763 | 0.083 . | 0.161 . | -0.521 | -0.527 | 0.237** | -0.055 |
| ***Ct* 2** | 0.072** | 0.022** | 0.086** | 0.254** | 0.261** | -0.216 | 0.027 . | 0.042 . | -0.241 | -0.127 | 0.179** | -0.039 |
| ***Ct* 3** | 0.039** | 0.012** | 0.047** | 0.111** | 0.140** | -0.090 | 0.013 . | 0.023 . | -0.117 . | -0.063 | 0.071* | -0.018 |
| ***Ct* 4** | 0.002** | -0.006 . | 0.003** | 0.019** | 0.011** | -0.007 | 0.001 . | 0.001 . | -0.089 | -0.005 | 0.006** | -0.001 |

### Model parameters for univariate analyses

The OU2VA model (via the *OUwie R* package; Beaulieu et al. 2012) was best-fitting among the univariate models for both the convergence simulations and divergence simulations (Table 2). This model is a multiple-regime OU model that allows the rate (σ) and attraction (α) parameters to vary between regimes (i.e., 'gliders' and 'non-gliders'), and these parameter values are reported in Table S2. The rates are considerably greater in the divergence simulations compared to the convergence simulations. For empirical convergence studies, we recommend close examination of parameter values. For example, especially fast rates for fitted OU models, such as those seen here for divergence simulations, may be an indicator of divergence rather than convergence. Compared to the rate parameter, the attraction parameter values do not show as much difference between convergence versus divergence simulations (Table S2).

**Table S2**. Rate (σ) and attraction (α) parameter values for the OU2VA model fitted to univariate data (PC1 scores). Values are means from 15 simulated datasets. Only the OU2VA results are provided because OU2VA was the best-fitting model in all analyses (Table 2). Note that rates are considerably greater in the divergence simulations compared to the convergence simulations.

| Simulations | Optimum | 'Gliders' | | 'Non-gliders' | |
|---|---|---|---|---|---|
| | | *σ* | α | *σ* | α |
| **Convergence** | 0 | 0.0369 | 0.0230 | 1.0118 | 0.0001 |
| | 20 | 0.0185 | 0.0193 | 1.0951 | 0.0005 |
| | 40 | 0.0093 | 0.0169 | 1.0109 | 0.0001 |
| | 60 | 0.0093 | 0.0131 | 1.0209 | 0.0003 |
| | 80 | 0.0114 | 0.0131 | 1.0092 | 0.0008 |
| **Divergence ('Constrained')** | 40 | 0.5584 | 0.0150 | 1.0432 | 0.0002 |
| | 60 | 0.0723 | 0.0149 | 1.0534 | 0.0009 |
| | 80 | 3.1667 | 0.0129 | 1.0573 | 0.0009 |

**Unconstrained divergence**

For our primary selection-based divergence simulations, we 'constrained' the focal lineages ('gliders') to one region of morphospace by limiting the OU-evolved divergent trait optima to be positive values, whereas all BM-evolved traits (of both 'gliders' and 'non-gliders') could be positive or negative (see Methods and Figure S1). We believe that the 'constrained divergence' simulations help to mimic empirical datasets in which lineages exhibit some morphological similarities but are still geometrically divergent.

Nonetheless, we also simulated 'unconstrained divergence' for a smaller subset of evolutionary scenarios (see discussion on subset analyses in the Methods) in which the 'glider' trait optimum values were not limited to positive values. The results of convergence measures applied to these simulations are provided in Table S3. The results indicate that all convergence measures correctly identify the 'gliders' to be divergent. For instance, the two-regime OU model (mvOU2) is out-performed by the single-regime mvBM1 and mvOU1 models, indicating a lack of evidence for 'glider' lineages converging on a shared adaptive peak. The mean *C*1 results are greater than zero, indicating a small amount of convergence. However, in contrast to *Ct*-measures, *C*-measures are not permitted to be less than zero (see Methods). Thus, the mean *C*1 results are biased toward being greater than zero because none of the 15 simulations can have values less than zero, and the small positive value is likely due to the occasional convergence of the ’glider’ lineages by chance.

**Table S3**. Convergence measures applied to ‘unconstrained divergence’ simulations in which focal taxa (‘gliders’) were permitted to evolve in any direction (see Methods and Figure S1). Results are the means of 15 simulated datasets for each trait optimum value. The Akaike weights are small-sample corrected Akaike weights. None of the results for the distance-based convergence measures are statistically significant. For θ results, we report $\theta_{real}$ standardized to phylogenetic distance between clades. See Tables 1 and 2 for more information.

| | | **Trait optimum of focal taxa** | | |
|---|---|---|---|---|
| | | **40** | **60** | **80** |
| **Model-ftting analsyes (Akaike weights)** | mvBM1 model | 1.000 | 1.000 | 1.000 |
| | mvOU1 model | 0.000 | 0.000 | 0.000 |
| | mvOU2 model | 0.000 | 0.000 | 0.000 |
| **Distance-based convergence measures** | θ (*RRphylo* ) | 0.509 | 0.468 | 0.530 |
| | Wheatsheaf index | 0.840 | 0.640 | 0.481 |
| | *C* 1 | 0.056 | 0.064 | 0.037 |
| | *Ct* 1 | -0.101 | -0.081 | -0.108 |

**Influence of ancestral state reconstructions**

As discussed in the main text, the assumption of a BM model of evolution for ancestral state reconstructions may often dilute *Ct* scores. We tested this assumption by comparing *Ct* scores obtained using our revised metric (with ancestral reconstructions via a BM model of evolution) to those obtained using a similar approach but employing the generated (‘true’) ancestral morphologies. The latter method used saved node values for each simulated morphological trait. Comparison between these two approaches was conducted using 15 simulated datasets, with four of six traits of ‘gliders’ OU-evolved and converging to optimum trait values of 0, 20, and 40 (see Fig 5), and the remaining two traits are BM-evolved. (These are the simulations used in the 'subset analyses' except that we excluded optima of 60 and 80.) Group-weighted *Ct* values showed consistently stronger convergence when true ancestral states were used in calculations (Fig. S8). These results indicate that the assumption that focal traits evolved via BM used to calculate *Ct* (and original *C*) scores can in some cases result in an underestimate of the degree to which lineages converge. *C* scores are also similarly influenced by this issue (Fig. S8).

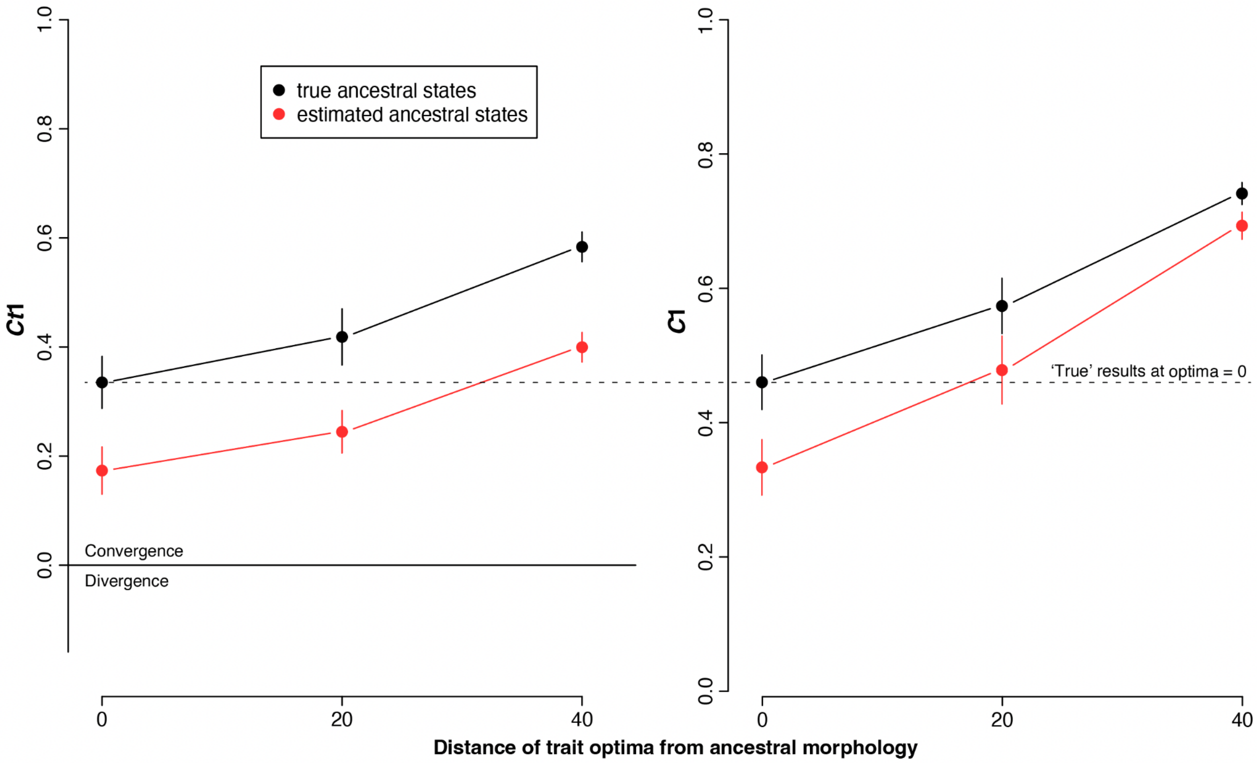


**Figure S8**. Comparisons of *Ct*1 (left) and *C*1 (right) scores obtained using ancestral reconstructions via a BM model of evolution ('red'; using the *calcConvCt* and *calcConv* functions) to those obtained using a similar approach but employing the generated ('true') ancestral morphologies ('black'). *Ct*1 scores and *C*1 scores are not directly comparable (except at complete convergence values of 1.0) because *C*1 tends to measure divergent taxa as convergent (e.g., Fig. 2B) and, unlike *C*1, *Ct*1 is not restricted to positive values. Nonetheless, we attempted to show a fairer comparison between the metrics by altering the y-axis scale so that the 'true' results of the two measures are aligned at optima of 0 (dashed line), which are results expected to have minimal biases related to morphospace position (e.g., see Figures 2A and S5). Note that *C*1 scores increase to a greater degree than Ct1 scores when optima are outliers, especially for the 'estimated ancestral state' analyses (see slopes of the red lines) – this likely reflects the issue highlighted in Figure 2A that *C*-measure values tend to be greater in morphological outliers.

To further examine the influence of the assumption of a BM mode of evolution during ancestral state reconstructions, we re-measured *Ct*1 scores using a range of ancestral reconstruction methods. These results are reported in Figure S9. In general, there was very little difference among *Ct*1 results, with results typically underestimating the true level of convergence, regardless of method used. Notably, none of the tested methods were close matches of the generating model, because they assume the same model of evolution for 'gliders' and 'non-gliders' (Figure S9). This is a limitation of most widely used methods for reconstructing ancestral states. Although users can in theory provide any manner of ancestral trait values to compute *Ct* metrics, we recommend a degree of caution. In empirical cases, where the true mode of evolution is unknown, treating putatively convergent groups separately from the rest of the sampled taxa risks circularity and may increase the chances of false positives.

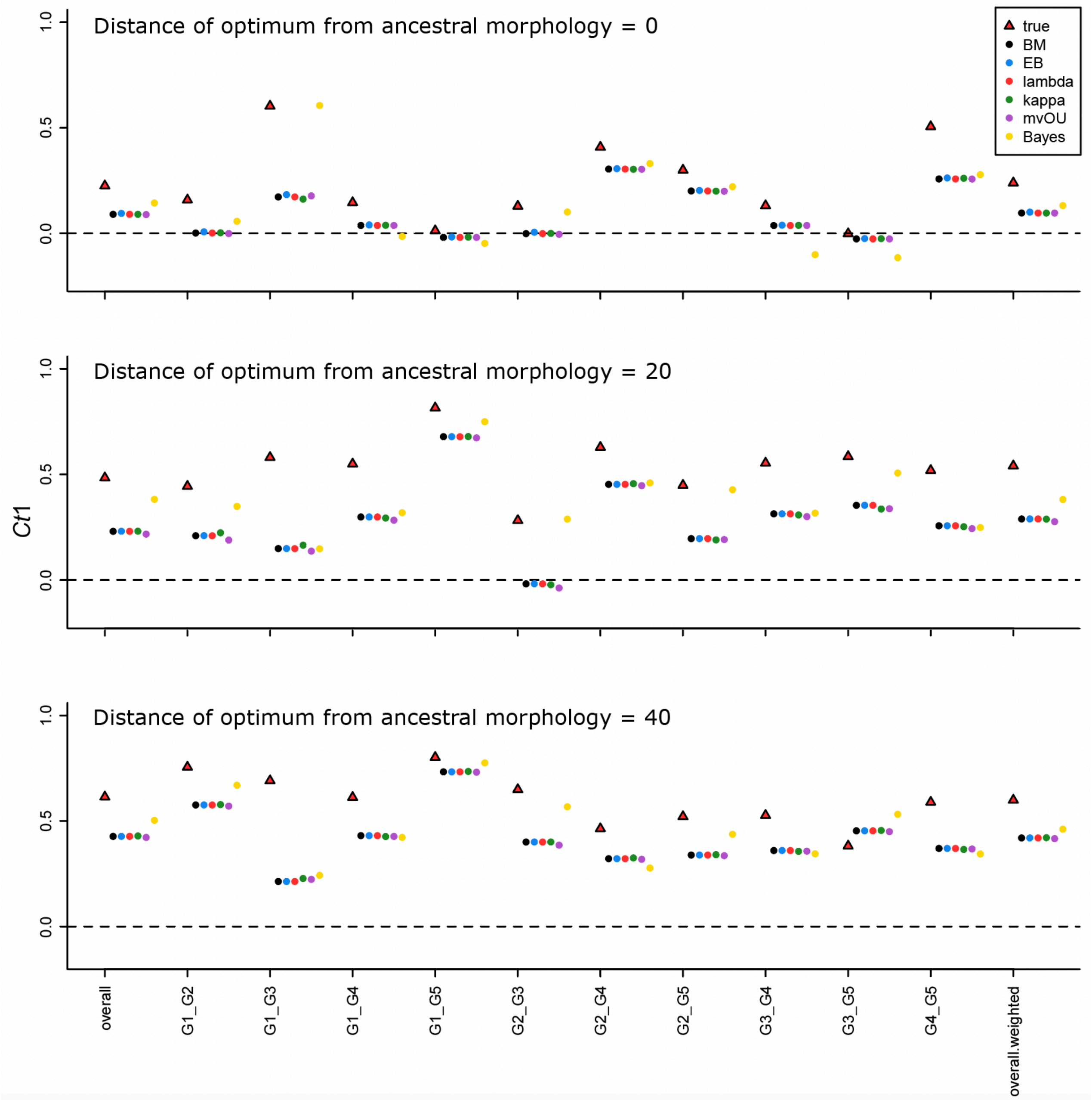


**Figure S9**. *Ct*1 values computed for simulated 'glider' lineages using a range of ancestral state reconstruction methods to calculate $D_{max.t}$, with results compared to *Ct*1 values calculated using the known ('true') ancestral state values for focal lineages. Each panel corresponds to a different simulated dataset where four of six traits within 'gliders' converged 0, 20, and 40 units from the morphological origin of the entire sample (see Methods). User supplied ancestral states for calculating *Ct*1 were obtained using the *phylopars R* package (Goolsby et al. 2017). These were computed assuming a single, universal Brownian Motion (BM), early burst (EB), lambda, kappa, or multivariate Ornstein-Uhlenbeck (mvOU) model. In addition, we reconstructed ancestral states using the *anc.Bayes* function in the *phytools R* package (Revell 2012) with four strict priors placed on the ancestral states at the base of each ‘glider’ clade. These were set to equal the values inferred for each trait by a BM model when all gliding lineages were dropped from the phylogeny. The effect was a more pronounced difference in ancestral states of gliding lineages and their immediate ancestors because observed states of modern gliders had reduced impact on the reconstructions of the latter. Setting such node limits is a simple way of mimicking variation across the tree in the evolutionary processes used to reconstruct ancestral states, and has some realistic basis (e.g., if fossil evidence can be used to constrain states at internal nodes). This approach achieved the results most closely resembling the true *Ct*1 values. However, in empirical cases, without the luxury of knowledge of the generating process, and in the absence of fossil evidence this approach should be adopted cautiously due to the risk of circularity. For each analysis, overall (weighted and unweighted; see Supplemental Methods) *Ct*1 values are given, along with each pairwise comparison between unique ‘glider’ groups (e.g., ‘G1_G2’ is the comparison between the first two ‘glider’ clades).

**LITERATURE CITED (in the Supplemental Information)**


Adams, D. C., and M. L. Collyer. 2018. Multivariate phylogenetic comparative methods: Evaluations, comparisons, and recommendations. Syst. Biol. 67:14-31.

Beaulieu, J. M., D. C. Jhwueng, C. Boettiger, and B. C. O’Meara. 2012. Modeling stabilizing selection: expanding the Ornstein–Uhlenbeck model of adaptive evolution. Evolution 66:2369–2383.

Felsenstein, J. 1985. Phylogenies and the comparative method. Am. Nat. 125:1–15.

Goolsby, E. W., J. Bruggeman, and C. Ané. 2017. Rphylopars: fast multivariate phylogenetic comparative methods for missing data and within-species variation. Methods Ecol. Evol. 8:22–27.

Huie, J. M., I. Prates, R. C. Bell, and K. de Queiroz. 2021. Convergent patterns of adaptive radiation between island and mainland *Anolis* lizards. Biol. J. Linn. Soc. Lond. 134:85–110.

Mahler, D. L., T. Ingram, L. J. Revell, and J. B. Losos. 2013. Exceptional convergence on the macroevolutionary landscape in island lizard radiations. Science 341:292–295.

Polly, P. D. 2019. Phylogenetics for Mathematica. Version 6.5. Department of Earth and Atmospheric Sciences, Indiana University: Bloomington, Indiana. https://pollylab.indiana.edu/software.html.

Revell, L. J. 2012. phytools: an R package for phylogenetic comparative biology (and other things). Methods Ecol. Evol. 3:217–223.

Stayton, C. T. 2015. The definition, recognition, and interpretation of convergent evolution, and two new measures for quantifying and assessing the significance of convergence. Evolution 69:2140–2153.

Stayton C. T. 2018. *Convevol: quantifies and assesses the significance of convergent evolution.* R package version 1.3. https:cran.r-project.org/package=convevol.

Upham, N., J. A. Esselstyn, and W. Jetz. 2019. Inferring the mammal tree: species-level sets of phylogenies for questions in ecology, evolution, and conservation. PLOS Biol. 17:e3000494.

Uyeda, J. C., D. S. Caetano, and M. W. Pennell. 2015. Comparative analysis of principal components can be misleading. Syst. Biol. 64:677–689.
